# Cohesin removal reprograms gene expression upon mitotic entry

**DOI:** 10.1101/678003

**Authors:** Carlos Perea-Resa, Leah Bury, Iain Cheeseman, Michael D. Blower

## Abstract

Entering mitosis, the genome is restructured to facilitate chromosome segregation, accompanied by dramatic changes in gene expression. However, the mechanisms that underlie mitotic transcriptional regulation are unclear. In contrast to transcribed genes, centromere regions retain transcriptionally active RNA Polymerase II (RNAPII) in mitosis. Here, we demonstrate that chromatin-bound cohesin is sufficient to retain RNAPII at centromeres while WAPL-mediated removal of cohesin during prophase is required for RNAPII dissociation from chromosome arms. Failure to remove cohesin from chromosome arms results in a failure to release elongating RNAPII and nascent transcripts from mitotic chromosomes and dramatically alters gene expression. We propose that prophase cohesin removal is the key step in reprogramming gene expression as cells transition from G2 to mitosis, and is temporally coupled with chromosome condensation to coordinate chromosome segregation with changes in gene expression.

**Highlights:**

- Mitotic centromere transcription requires cohesin
- Cohesin removal releases elongating RNA Pol II and nascent RNA from chromatin
- The prophase pathway reprograms gene expression during mitosis

## Introduction

During mitosis, interphase chromatin organization is erased as chromosomes are condensed and individualized in preparation for their segregation (Gibcus et al., 2018). As chromosomes condense, the pattern of gene expression is dramatically altered (Liang et al., 2015; Palazzo et al., 2007; Prescott and Bender, 1962). Mitotic transcriptional silencing is thought to occur by preventing new initiation through phosphorylation of the general transcription factors, TFIID and TFIIH (Akoulitchev and Reinberg, 1998; Segil et al., 1996), as well as phosphorylation of the C-terminal domain (CTD) of RNA Polymerase II (RNAPII) (Bellier et al., 1997). Additionally, p-TEF-b is activated at mitotic onset to promote the transcriptional run-off of paused RNAPII during prophase (Liang et al., 2015). In contrast to the bulk of the genome, centromeres escape general mitotic transcriptional inhibition and retain elongating RNAPII (Chan et al., 2012). The previously described mechanisms for mitotic transcriptional silencing cannot account for the persistence of active RNAPII at centromeres, suggesting that additional mechanisms regulate the spatial pattern of mitotic transcriptional inhibition.

Cohesin is a major component of interphase and mitotic chromosomes regulating transcription, DNA double-strand break repair or sister chromatid cohesion (Nasmyth and Haering, 2009). Cohesin is linked to transcription through its interaction with Mediator, which promotes enhancer-promoter interactions (Kagey et al., 2010), the formation of topologically-associated domains (TADs) (Rao et al., 2017), interaction with the super-elongation complex (Izumi et al., 2015), and recruitment of transcriptional activators to transcription factor hotspots (Yan et al., 2013). In interphase cells, cohesin is widely distributed throughout the genome where it regulates both chromosome structure and cohesion between replicated chromatids. During mitotic prophase, the kinases Plk1, Aurora B, and Cdk1 phosphorylate the cohesin-associated proteins Sororin and SA2 to promote cohesin removal from chromosome arms by the cohesin release factor WAPL, in a process termed the ‘prophase pathway’ (Haarhuis et al., 2014). Centromeric cohesin is protected from WAPL-mediated removal by Shugoshin proteins until the metaphase-anaphase transition when cohesin rings are cleaved by the Separase protease (Uhlmann et al., 2000). While it is established that cohesin loops formed in *cis* regulate chromatin architecture and gene expression during interphase, it is not clear if the cohesin tethering duplicated sister chromatid also regulates gene expression during mitosis.

Here, we use centromere transcription as a model to study the mechanisms of mitotic transcription inhibition. We find that dissociation of RNAPII and cohesin from mitotic chromosomes are temporally and spatially correlated. We show that chromatin-localized cohesin is both necessary and sufficient to retain elongating RNAPII on mitotic chromosomes. Additionally, we demonstrate that static cohesin prevents dissociation of elongating RNPII from both interphase and mitotic chromosomes. Our results indicate that WAPL-mediated removal of cohesin from chromosomes during prophase functions to regulate gene expression and that mitotic centromere transcription is unlikely to play an active role in mitosis.

## Results

### Cohesin promotes selective retention of RNA Pol II at centromeres in mitosis

The differential localization of RNAPII during mitosis could be explained by its active recruitment to centromeres or reflect selective protection from a global removal pathway (Fig. 1A). To determine how RNAPII is selectively localized to centromeres during mitosis, we analyzed the distribution of actively-elongating (CTD phospho-Ser 2, pS2) and promoter-paused (CTD phospho-Ser 5, pS5) RNAPII during early stages of mitosis using non-transformed human RPE-1-hTERT cells (Fig. 1B-C). RNAPII pS2 localized to mitotic centromeres from G2 through metaphase, but then became significantly reduced at centromeres in anaphase as sister chromatids separated (Fig. 1B). In contrast, CTD pS5 intensity steadily declined throughout mitosis (Fig. 1C). We observed similar results for RNAPII localization in *Xenopus* egg extracts (Fig. S1). To understand the differential localization of RNAPII pS2 to mitotic centromeres, we analyzed both the absolute fluorescent intensity at centromeres and its behavior relative to chromosome arms. We found that the absolute levels of RNAPII pS2 at centromeres remained constant from G2 to metaphase (Fig. 1B). Similarly, chromatin immunoprecipitation (ChIP) analysis using RNAPII CTD pS2 antibodies revealed that RNAPII is present at human centromeres at similar levels in G2 and metaphase, but is removed from the β-actin gene during mitosis (Fig. 1D). These results suggest that the relative enrichment of RNAPII at mitotic centromeres occurs through its selective removal from chromosome arms.

**Figure 1.**
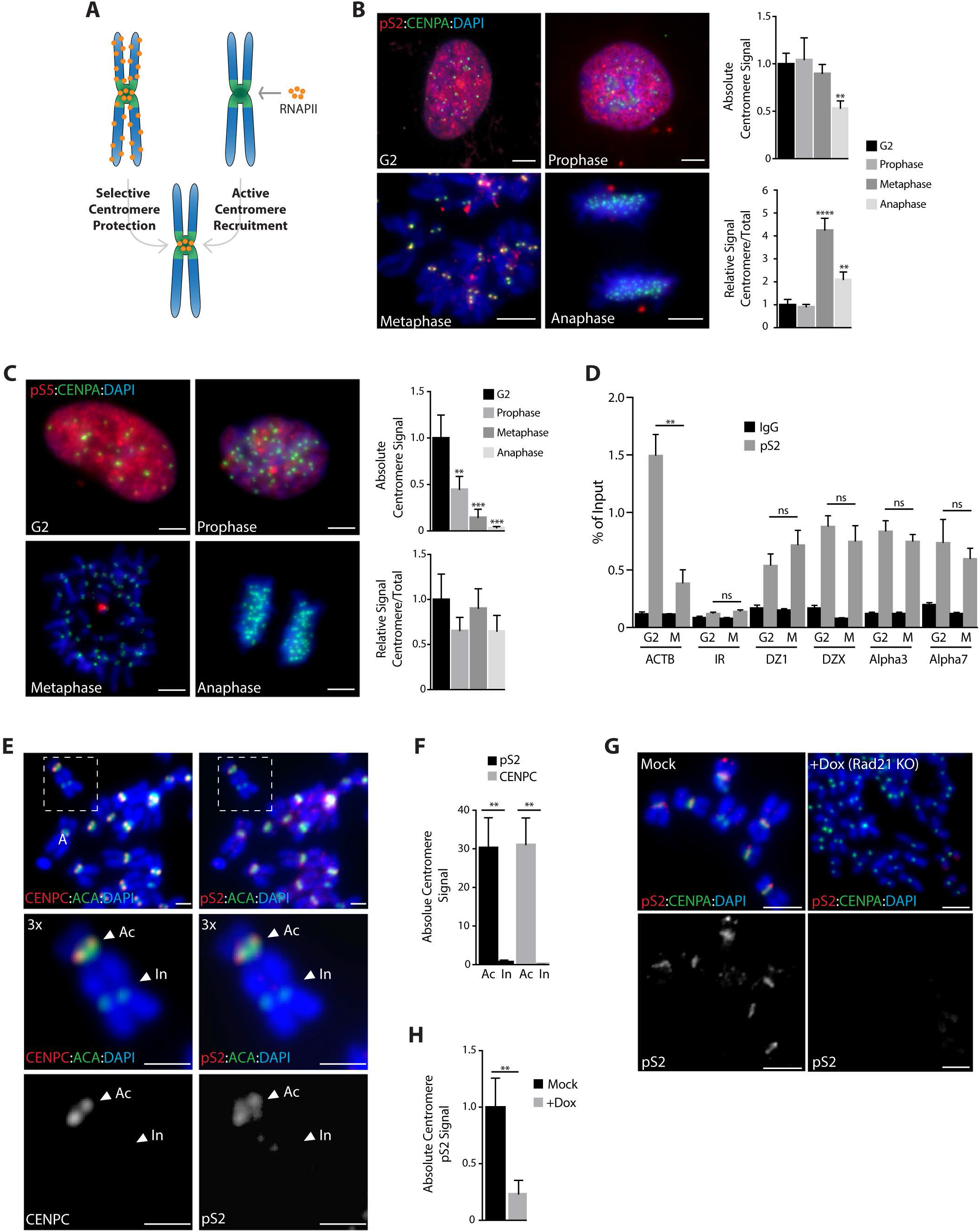
Cohesin promotes the selective retention of RNA Pol II at centromeres in mitosis. **A.** Either selective chromosome arm removal or active centromere recruitment early in prophase could explain RNA Pol II enrichment at metaphase centromeres. **B-C** pS2 and pS5 localization (red) in chromosome spreads from RPE-1 cells cycled from interphase (G2) to anaphase. Anti-CENPA antibodies (green) and DAPI (blue) were used to visualize centromeres and DNA respectively. Scale bars, 10 μm. Right, quantification of absolute and relative centromere signal. **, P < 0.005; ***, P < 0.001 (one-way ANOVA, Dunnett’s test). **D** pS2 occupancy at centromeres of chromosomes 1, 3, 7 and X, represented as the % of input recovered following ChIP in G2 or metaphase (M) RPE-1 cells. Occupancy values at the ACTB gene and at a random intergenic region (IR) were included as positive and negative controls, respectively. **, P < 0.005; (ns) no significant differences (t-test). **E-F** Analysis of CENPC and pS2 localization (red) in metaphase spreads from dicentric MDA-MB 435 cells. ACA (anti-centromere antibodies; green) stains both active (Ac) and inactive (In) centromeres while CENPC and pS2 (red) are only recruited to the active kinetochore region. Scale bars, 5 μm. Quantification of pS2 and CENPC centromere signal is plotted on F. **G-H** Analysis of pS2 localization (red) on metaphase spreads from doxycycline inducible Rad21 KO cells. Anti-CENPA (green) and DAPI (blue) were used to visualize centromeres and DNA, respectively. Scale bars, 10 μm. Quantification of absolute centromere levels is represented in H. **, P < 0.005 (t-test). In all cases error bars indicate the standard deviation of the mean.

In vertebrate cells, centromeres are specified epigenetically by the presence of histone variants, and are not defined by precise DNA sequences. To determine if the retention of RNAPII on mitotic chromosomes is linked to the active centromere or is a general property of a-satellite DNA, we analyzed RNAPII localization on a dicentric human chromosome. Structurally dicentric chromosomes contain two a-satellite arrays, but form an active centromere on only one array. Consistent with previous results (Chan et al., 2012), RNAPII only localized to the kinetochore-forming array of a dicentric chromosome (Fig. 1E-F). Thus, RNAPII retention is linked to functional centromere activity. In addition to displaying differences in kinetochore formation, the inactive centromere on dicentric chromosomes was notably separated from its sister partner indicating the absence of cohesion. To evaluate the role of sister chromatid cohesion in retaining RNAPII at centromeres, we depleted the cohesin subunit Rad21 (Fig. S2). Strikingly, Rad21 depletion resulted in a complete loss of RNAPII localization to mitotic chromosomes (Fig. 1G-H) and in reduced levels of centromeric RNAs (Fig. S2C), suggesting that the presence of cohesin helps retain RNAPII at centromeres. Consistent with a close connection between cohesin and RNAPII, we found that the spatial pattern of cohesin removal during prophase closely parallels that of RNAPII from chromosome arms in both human cells and *Xenopus* egg extracts (Fig. S3).

### Ectopic cohesin retains active RNA Pol II on metaphase chromosomes

To determine if prophase cohesin removal is required for the eviction of RNAPII from mitotic chromosomes, we generated an Auxin Inducible Degron (AID) human cell line for the cohesin release factor WAPL (Fig. S4A-B). Addition of auxin (IAA) to WAPL-AID cells resulted in WAPL degradation, cohesin retention and the persistence of sister chromatid cohesion along the entire length of mitotic chromosomes (Fig. S4C-G). Strikingly, WAPL degradation following auxin addition also resulted in the retention of RNAPII pS2 on the arms of mitotic chromosomes while no effect was detected on pS5 localization (Fig. 2A-C; Fig. S4H-I). Similar results were obtained following WAPL RNAi in human cells or WAPL immuno-depletion in *Xenopus* egg extracts (Fig. S5). In addition, preventing cohesin removal by inhibition of either Aurora B or Plk1 also resulted in retention of elongating RNAPII (pS2) on mitotic chromosome arms in both human cells and *Xenopus* egg extracts (Fig. S6). Retention of RNAPII pS2 in WAPL-AID cells was dependent on the presence of cohesin as co-depletion of WAPL and Rad21 resulted in a lack of RNAPII pS2 on mitotic chromosomes, similar to the phenotype observed after Rad21 depletion (Fig. S7). Taken together, our data suggest that cohesin removal from chromosome arms during mitotic prophase is the key step in the eviction of RNAPII from chromosomes during mitosis. Cohesin protection at active centromeres explains the selective retention of RNAPII at centromeres during mitosis.

**Figure 2.**
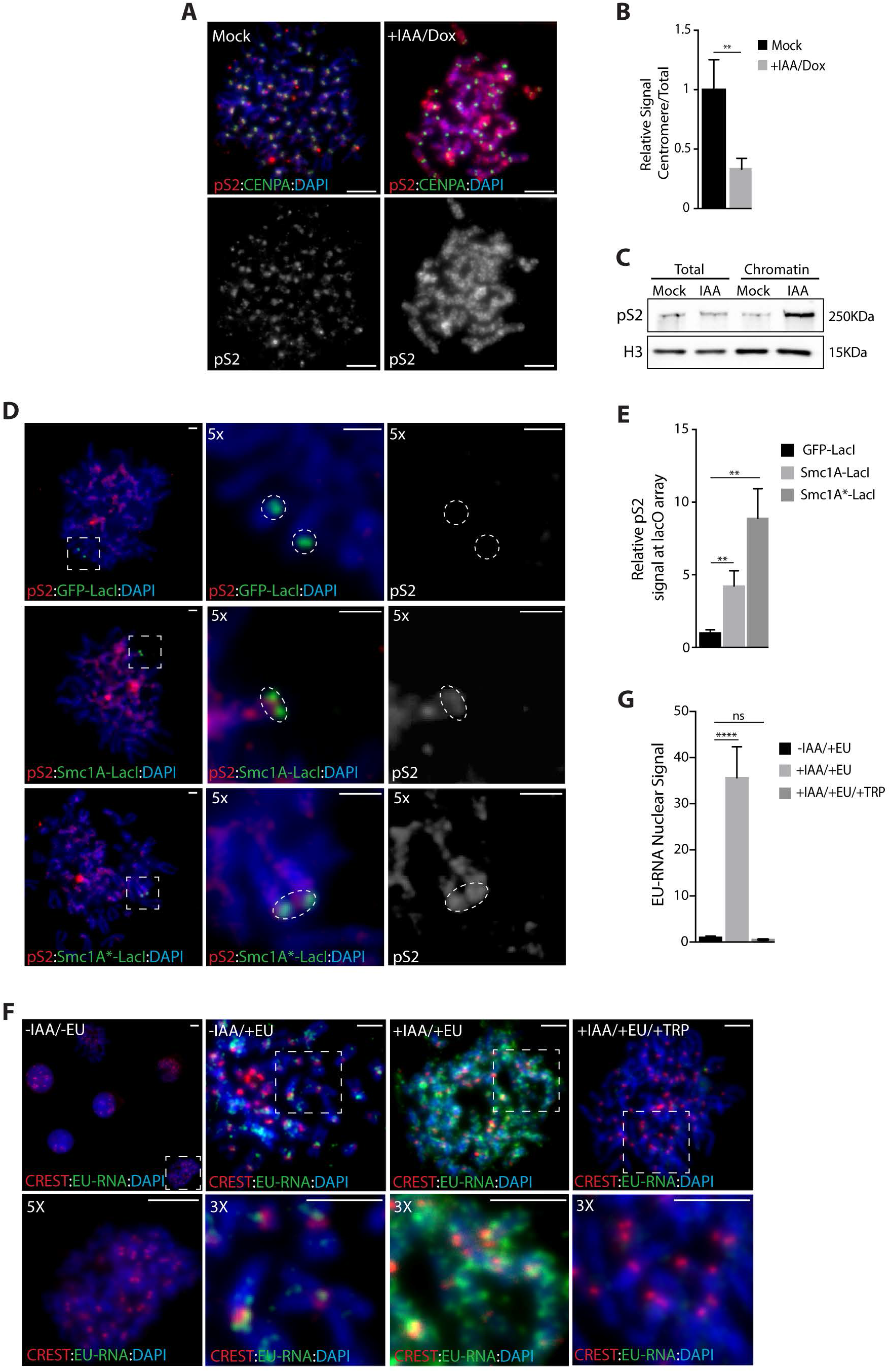
Ectopic cohesin retains RNA Pol II activity on metaphase chromosomes. **A** Analysis of pS2 localization (red) in metaphase spreads from AID-WAPL cells. Cells were incubated with DMSO (Mock) or IAA and doxycycline for 48 h to promote AID-WAPL degradation. Anti-CENPA antibodies (green) and DAPI (blue) were used to visualize centromeres and DNA, respectively. Scale bars, 5 μm. **B** Quantification of pS2 centromere enrichment in images from A. Data represent relative enrichment as a ratio of centromere to total nuclear signal. Levels were normalized to control cells. **C** pS2 levels in total or chromatin protein fractions from metaphase AID-WAPL cells measured by Western blotting. Levels of histone H3 provide a loading control. **D** pS2 localization (red) in metaphase spreads from U2OS cells following the tethering of LacI-tagged proteins to the lacO locus. GFP-LacI, SMC1A(WT)-LacI, or SMC1A*(mt)-LacI fusion proteins were transiently expressed and metaphase chromosome spreads were analyzed 48 h after transfection. Anti-GFP or anti-FLAG antibodies were used to detect the LacI-tagged proteins (green) and DAPI staining to visualize chromosomes (blue). Scale bars, 2 μm. **E** Quantification of pS2 levels at the lacO array following cell transfection as detailed in D. **F-G** Analysis of nascent RNA localization on metaphase chromosome spreads from AID-WAPL cells. Cells were incubated in the presence or absence of EU for nascent RNA labeling. EU incorporation was detected by Click-iT chemistry (green). Anti-centromere antibodies (CREST; red) and DAPI staining (blue) were used for centromere and chromosome labelling, respectively. Scale bars, 5 μm. **G** Quantification of total EU nuclear fluorescence. Data represent the mean of three independent experiments and values were normalized to control cells (-IAA/+EU). In all cases, error bars indicate the standard deviation of the mean. **, P < 0.005; ****, P < 0.0001 (t-test).

To determine if cohesin is sufficient to promote transcription of an ectopic locus during mitosis, we tethered either wild type Smc1a (a subunit of cohesin) or an ATPase-deficient form of Smc1a that is resistant to WAPL-mediated removal (Elbatsh et al., 2016) to an array of lac operator sequences located at the arm of chromosome 1 in U2OS cells (Janicki et al., 2004). Tethering of Smc1a-LacI led to the recruitment of the cohesin subunit Smc3. ~50% of the cohesin signal was retained on chromatin following treatment with IPTG to disrupt the LacI-lacO interaction, suggesting that the entire cohesin complex is recruited to DNA as a ring encircling chromatin (Fig. S8A-D). Tethering of Smc1-LacI, but not GFP-LacI, resulted in strong recruitment of elongating RNAPII (pS2) to interphase and mitotic chromosomes (Fig. 2D-E; Fig. S8E-F). During mitosis, RNAPII recruitment was more efficient when WAPL-resistant cohesin was tethered to chromatin (Fig. 2E). Taken together, our results demonstrate that cohesin removal from euchromatin by the ‘prophase pathway’ helps evict RNAPII from mitotic chromosomes and that retention of cohesion is sufficient to allow RNAPII to persist on chromosomes.

To understand if cohesin retention on mitotic chromosomes results in enzymatically active RNAPII, we next pulse-labeled control and WAPL-AID cells with the RNA label EU during mitosis (Jao and Salic, 2008). Mitotic chromosome spreads from control cells displayed newly-synthesized RNA present primarily at centromeres (Fig. 2F). In contrast, WAPL-AID cells displayed newly-synthesized RNA throughout the chromosomes (Fig. 2F-G). Importantly, this nascent-RNA signal on mitotic chromosomes in WAPL-AID cells was eliminated by treatment with triptolide, an inhibitor of RNAPII transcription initiation (Titov et al., 2011) (Fig. 2F-G), likely through RNAPII degradation following extended triptolide treatment (Novais-Cruz et al., 2018).

### Cohesin retention alters gene expression reprogramming during the G2-M transition

Failure to remove cohesin from euchromatin in prophase in WAPL-depleted cells results in a broad distribution of active RNAPII on mitotic chromosomes. To understand the impact of cohesin retention on the transcriptome, we performed RNA sequencing of newly synthesized transcripts (EU labeled) from G2 and mitotically-arrested control and WAPL-depleted cells. Principle Component Analysis (PCA) indicated that experimental replicates clustered and that G2 and mitotic samples were well separated by PC1 in control cells (Fig. 3A). WAPL-depleted cells displayed significantly different gene expression from control cells, consistent with previous work (Haarhuis et al., 2017; Tedeschi et al., 2013). In WAPL-depleted cells, G2 and mitotic cells were not well-separated by PC1, suggesting a similar pattern of gene expression persists between G2 and mitosis. Analysis of the sequence coverage across all genes demonstrated a significant peak at the 5’ end of genes in G2 cells (Fig. 3B, S9), likely representing paused, promoter-proximal RNAPII. Upon entry into mitosis, this 5’ peak was lost in both control and WAPL-depleted cells, suggesting that persistent cohesin does not affect the process of RNAPII activation by p-TEF-b to release from its promoter-proximal pause state. Indeed, we found that inhibition of several different steps of transcriptional initiation during mitosis did not affect the distribution of RNAPII on mitotic chromosomes in control or WAPL-depleted cells (Fig. S10). This suggests that persistent cohesin does not promote transcription initiation during mitosis.

**Figure 3.**
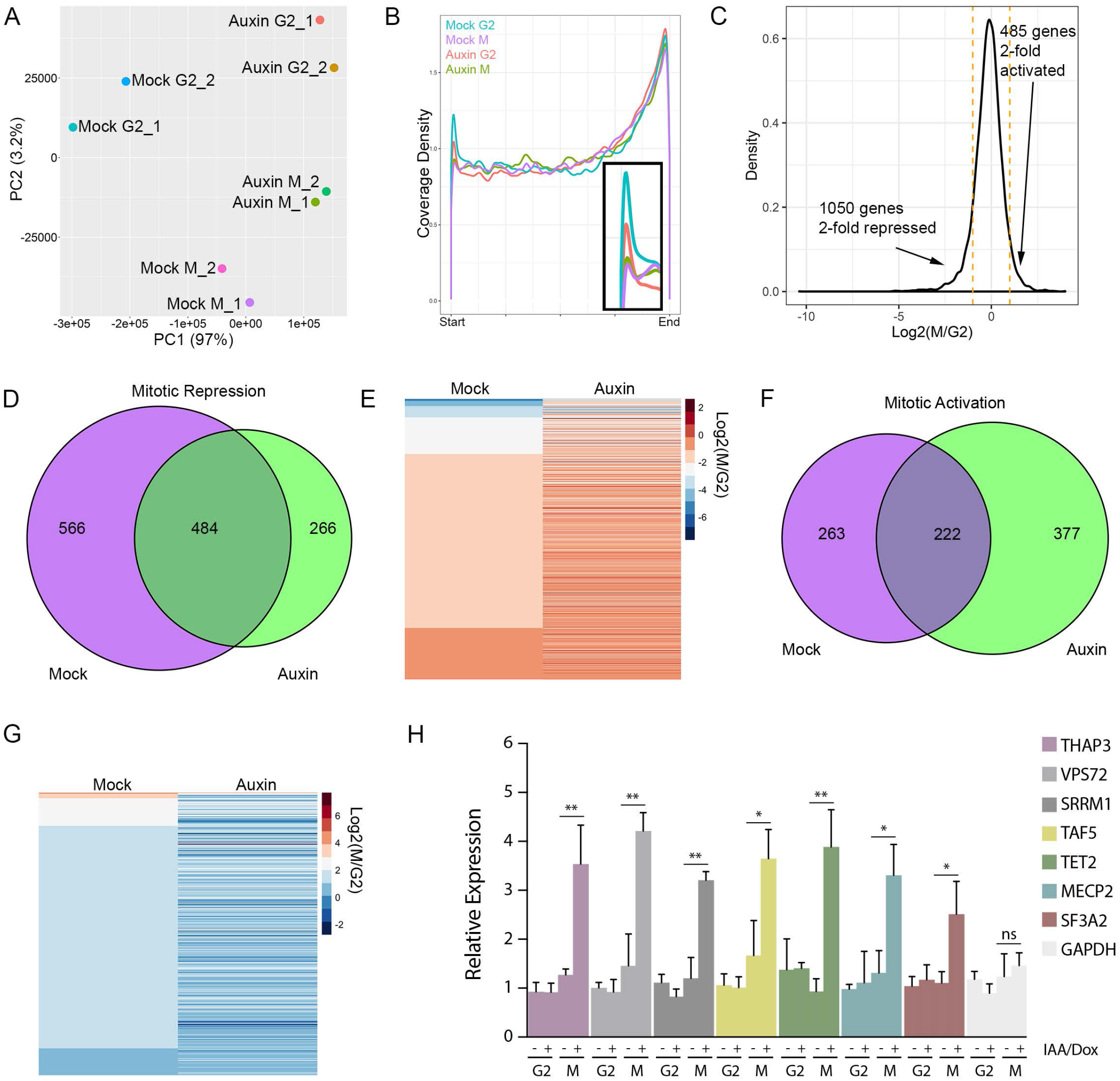
Cohesin retention alters gene expression reprogramming in the G2-M transition. **A** Principle Component Analysis (PCA) of gene expression in G2 and mitosis from replicate cultures of mock or auxin-treated WAPL-AID cells. **B** Metaplot of relative sequencing coverage across all genes expressed at FPKM > 10. **C** Histogram of changes in gene expression between G2 and mitosis in control cells. **D** Venn diagram of overlap of mitotically-repressed genes in control and auxin-treated WAPL-AID cells. **E** Heatmap of mitotically repressed genes in control and auxin-treated WAPL-AID cells. **F** Venn diagram of mitotically activated genes in control and WAPL-AID cells. **G** Heatmap of mitotically activated genes in control and auxin-treated WAPL-AID cells. **H** Q-RT-PCR validation of EU-labeled transcription in WAPL-AID cells in G2 and mitosis. Error bars represent the SD of the mean. *, P < 0.05; **, P < 0.005 (t-test). (ns) no significant differences.

In control cells, we found that hundreds of genes were both transcriptionally activated and repressed during mitosis (Fig. 3C), consistent with previous work (Liang et al., 2015). However, of the 1050 genes that were > 2-fold repressed in control cells, less than half of these genes were repressed in WAPL-depleted cells (Fig. 3C-E, H, S9). Thus, persistent cohesin results in decreased mitotic transcriptional repression. Reciprocally, of the 485 genes that were > 2-fold transcriptionally activated in control cells, less than half of these genes were activated in WAPL-depleted cells (Fig. 3C, F-G). In addition to transcription of coding regions, we observed that expression of intergenic enhancers both increased and decreased during mitosis. However, WAPL depletion did not have a global effect on enhancer expression (Fig. S11). Similarly, RNAs mapping to the centromere did not change between G2 and mitosis in control cells and were unaffected by WAPL depletion (Fig. S12). Taken together, these results suggest that the ability to remove cohesin from chromosomes is important for the cell cycle dynamics of RNAPII and that persistent cohesin results in dampened transcriptional changes between G2 and mitosis.

### Cohesin retention impairs the release of elongating RNA Pol II from chromosomes

To determine if persistent cohesin affects the dynamics of RNAPII association with chromosomes, we performed fluorescent recovery after photobleaching (FRAP) analysis in GFP-RPB1 knock-in cells (Steurer et al., 2018). In control cells during interphase, the fluorescent recovery of GFP-RPB1 fit to a three-component model representing the different chromatin-bound RNAPII populations during transcription: initiation (A, k1), promoter-paused (B, k2), and elongating states (C, k3) (Fig. 4 A-B and Fig. S13A-B), consistent with prior work (Darzacq et al., 2007; Steurer et al., 2018). Diffusion did not contribute to RNAPII FRAP recovery, suggesting that most RNAPII is chromatin-bound (Fig. S14B-E)(Darzacq et al., 2007; Steurer et al., 2018). In contrast to controls, WAPL-depleted interphase cells showed significantly slower fluorescent recovery of GFP-RPB1. Modeling of this curve behavior indicated that this difference was primarily due to a higher fraction of elongating RNAPII, 44% vs 51%, and a ~ 35 % lower dissociation rate of elongating RNAPII (k3) (Fig. 4A-B, Fig. S13B). As a consequence, the pool of RNAPII available for promoter binding is reduced as suggested by a lower pseudo-on rate (A) (Fig. 4B, Fig. S13C). During mitosis, control cells displayed a rapid fluorescent recovery of GFP-RPB1. Analysis of the turnover dynamics indicated that a two-component model could explain the fast fluorescence turnover primarily based on the presence of freely diffusing (non-chromatin bound) RNAPII (Fig. 4C-D, Fig. S14F). In contrast, WAPL-depleted cells displayed a much slower, diffusion-independent recovery with a behavior resembling that of interphase, elongating RNAPII dynamics (Fig. 4C-D, Fig. S14G). In summary, analysis of GFP-RBP1 fluorescent recovery demonstrates that cohesin removal promotes the release of elongating RNAPII from chromatin in interphase and mitotic cells.

**Figure 4.**
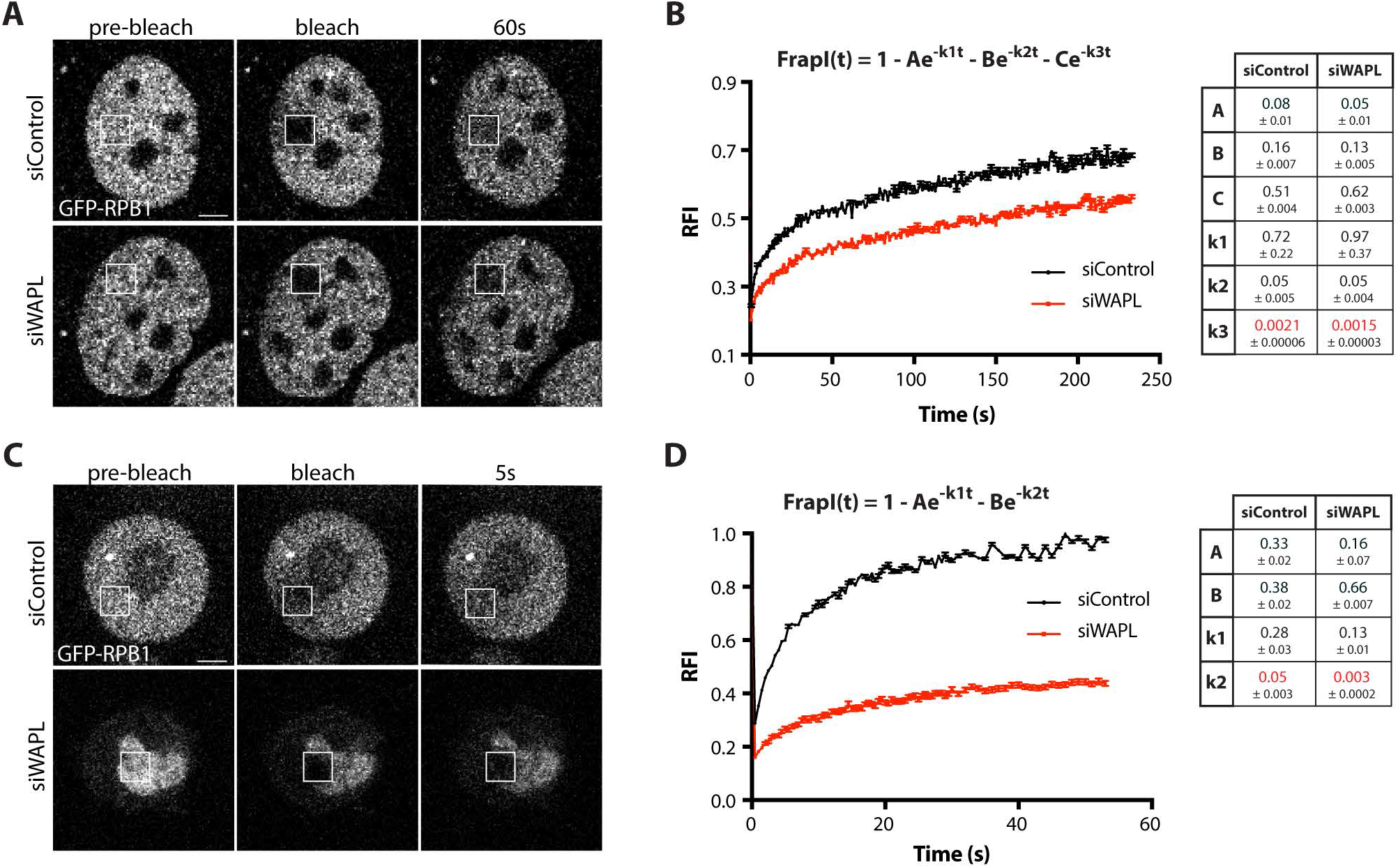
Cohesin retention impairs elongating RNA Pol II release from chromosomes. GFP-RPB1 knock-in fibroblasts were used for the analysis of RNA Pol dynamics by FRAP. Cells were bleached 48 h after transfection with either siControl or siWAPL siRNAs followed by imaging for fluorescence recovery. Images of the indicated times are shown for representative cells in interphase (A) and mitosis (C). Scale bars, 3 μm. **B, D** Relative fluorescence intensity (RFI) between bleached (dash line squares) and non-bleached reference areas are plotted in time. Measurements from interphase and mitotic cells were fitted to three- or two-component models, respectively. The equation and a table containing the values estimated for each parameter are shown to the right. n ≥ 14 cells were used for each condition from two independent experiments. In all cases, error bars indicate the standard error of the mean (sem).

## Discussion

### Role of cohesin dynamics on gene expression reprogramming

During mitotic entry interphase chromatin structure is erased and gene expression is dramatically altered (Gibcus et al., 2018; Liang et al., 2015; Palozola et al., 2017) yet the molecular mechanisms controlling this process are not clear. Our results demonstrate that the prophase pathway of cohesin removal is a key factor in rewiring transcription upon mitotic entry. Removal of cohesin from chromosome arms facilitates the release of elongating RNAPII while cohesin retention at centromeres explains how transcription persist at centromeres of metaphase chromosomes. As cells exit mitosis transcription is rapidly reactivated (Palozola et al., 2017; Teves et al., 2018) in a manner that is temporally correlated with cohesin loading, yet it is not clear if cohesin is required for gene reactivation following mitosis. During prophase WAPL-mediated release of cohesin from chromosome arms safeguards cohesin from Separase decay (Kueng et al., 2006), which may facilitate the rapid deposition of cohesin on chromosomes later in mitosis. This model is supported by the localization of cohesin loaders Scc2 (Nipbl) and Scc4 to telophase chromosomes (Watrin et al., 2006). Whether cohesin loading in late telophase is required for gene expression re-activation early in G1 will require further studies. Interestingly, cohesin remains bound to binding sites of transcription factors (TF) clusters in mitosis while TFs are evicted, suggesting a role for cohesin in cell memory (Yan et al., 2013). In addition to mitosis, gene expression shows an extensive reprogramming following cell differentiation during development. Mutations on genes encoding the cohesin subunits SMC1A, SMC3 and RAD21, or their regulators NIPBL and HDAC8, have been identified in the Cornelia de Lange syndrome (CdLS), a cohesinopathy causing severe developmental disorders (Banerji et al., 2017). Importantly, cells from CdLS patients exhibit no cohesion defects but significantly altered transcriptional profiles (Mannini et al., 2015), supporting a central role for cohesin and its regulation rewiring gene expression during development.

### Mitotic centromere transcription

Retention of elongating RNAPII at centromeres has been observed in metaphase chromosomes of human, fly and frog cells(Blower, 2016; Chan et al., 2012; Molina et al., 2016; Rosic et al., 2014), and several roles for mitotic centromere transcription have been proposed (Blower, 2016; Chan et al., 2012; Liu et al., 2015). However, a recent study has questioned the role of mitotic transcription for accurate chromosome segregation (Novais-Cruz et al., 2018). Our results demonstrate that elongating RNAPII is passively retained at centromeres in mitosis as a byproduct of cohesin retention. In WAPL-depleted cells, active RNAPII pS2 is retained on chromosome arms as a consequence of failure in cohesin release. Previous studies have shown modest chromosome segregation defects following WAPL-depletion as a consequence of failure in Aurora B (AurB) and Sugoshin 1 (Sgo1) concentration at metaphase centromeres (Haarhuis et al., 2013; Kueng et al., 2006; Tedeschi et al., 2013). Interestingly, both AurB and Sgo1 proteins have RNA-binding activity (Jambhekar et al., 2014; Liu et al., 2015). It will be important to determine if delocalization of Aurora-B and Sgo1 in WAPL-depleted cells results from ectopic transcription or cohesin retention. However, WAPL knockout cells are viable (Haarhuis et al., 2017) and exhibit modest chromosome segregation defects (Haarhuis et al., 2013; Tedeschi et al., 2013). Additionally, the ‘prophase pathway’ is not conserved in budding yeast (Haarhuis et al., 2014), suggesting that restricting transcription to the centromere during early mitosis is unlikely to be necessary for accurate chromosome segregation. We speculate that mitotic centromere transcription is a byproduct of the role of cohesin in regulating gene expression during interphase and is not required for accurate chromosome segregation.

### Regulation of transcription by cohesin in mitosis

Cohesin is a major constituent of genome architecture and regulates many aspects of gene transcription during interphase. Global analyses of cohesin distribution by ChIP-seq have demonstrated a correlation between the cohesin complex and active transcription (Busslinger et al., 2017; Schaaf et al., 2013). However, during early mitosis most cohesin and cohesin-mediated TADs are removed and it was not clear if chromatin-bound cohesin also regulated transcription at this cell stage. Our results reveal cohesin as the key factor determining the distribution of elongating RNA Pol II (pS2) on mitotic chromosomes. Consistent with previous studies (Chan et al., 2012), we find specific retention of active elongating pS2 while promoter paused RNA Pol II (pS5) is largely absent from metaphase chromosomes. Additionally, mitotic inhibition of transcription initiation does not affect the localization of RNAPII pS2 to centromeres and only elongating RNAPII is retained on mitotic chromosomes in WAPL-depleted cells. Collectively, these data support the absence of transcription initiation during mitosis limiting the role of cohesin to latter steps of transcription regulation. Interestingly, several studies suggest cohesin facilitates the transition of RNA Pol II from promoter-pause to the actively elongating state (Izumi et al., 2015; Schaaf et al., 2013). In addition, cohesin promotes transcription termination in yeast (Gullerova and Proudfoot, 2008). Our FRAP results reveal increased levels of chromatin-bound elongating RNAPII when cohesion is retained on mitotic chromosomes. We also observe a similar effect in interphase cells, but the effect is more subtle because transcription initiation is active during this cell stage. Thus, our work indicates that cohesin removal promotes release of elongating RNAPII from mitotic chromatin.

Here, we demonstrated that the removal of cohesin from chromosomes during prophase facilitates mitotic RNAPII dynamics and contributes to transcription inhibition during mitosis. Our results demonstrate that cohesin removal is coupled to transcriptional remodeling as a component of the process that restructures the genome in preparation for mitotic chromosome segregation. Our work suggests that the ‘prophase pathway’ of cohesin removal acts primarily to reprogram gene expression. Indeed, chromosome segregation can proceed in the absence of the prophase cohesin removal pathway (Gandhi et al., 2006; Kueng et al., 2006). Thus, the prophase pathway serves to counteract the interphase transcriptional functions of cohesin allowing the rewiring of the transcriptional program during mitosis.

## References

Akoulitchev, S., and Reinberg, D. (1998). The molecular mechanism of mitotic inhibition of TFIIH is mediated by phosphorylation of CDK7. Genes & development 12, 3541–3550.

Banerji, R., Skibbens, R.V., and Iovine, M.K. (2017). How many roads lead to cohesinopathies? Dev Dyn 246, 881–888.

Bellier, S., Dubois, M.F., Nishida, E., Almouzni, G., and Bensaude, O. (1997). Phosphorylation of the RNA polymerase II largest subunit during Xenopus laevis oocyte maturation. Molecular and cellular biology 17, 1434–1440.

Blower, M.D. (2016). Centromeric Transcription Regulates Aurora-B Localization and Activation. Cell reports.

Busslinger, G.A., Stocsits, R.R., van der Lelij, P., Axelsson, E., Tedeschi, A., Galjart, N., and Peters, J.M. (2017). Cohesin is positioned in mammalian genomes by transcription, CTCF and Wapl. Nature 544, 503–507.

Chan, F.L., Marshall, O.J., Saffery, R., Kim, B.W., Earle, E., Choo, K.H., and Wong, L.H. (2012). Active transcription and essential role of RNA polymerase II at the centromere during mitosis. Proceedings of the National Academy of Sciences of the United States of America 109, 1979–1984.

Darzacq, X., Shav-Tal, Y., de Turris, V., Brody, Y., Shenoy, S.M., Phair, R.D., and Singer, R.H. (2007). In vivo dynamics of RNA polymerase II transcription. Nature structural & molecular biology 14, 796–806.

Elbatsh, A.M.O., Haarhuis, J.H.I., Petela, N., Chapard, C., Fish, A., Celie, P.H., Stadnik, M., Ristic, D., Wyman, C., Medema, R.H., et al. (2016). Cohesin Releases DNA through Asymmetric ATPase-Driven Ring Opening. Molecular cell 61, 575–588.

Fishilevich, S., Nudel, R., Rappaport, N., Hadar, R., Plaschkes, I., Iny Stein, T., Rosen, N., Kohn, A., Twik, M., Safran, M., et al. (2017). GeneHancer: genome-wide integration of enhancers and target genes in GeneCards. Database (Oxford) 2017.

Gandhi, R., Gillespie, P.J., and Hirano, T. (2006). Human Wapl is a cohesin-binding protein that promotes sister-chromatid resolution in mitotic prophase. Current biology: CB 16, 2406–2417.

Gibcus, J.H., Samejima, K., Goloborodko, A., Samejima, I., Naumova, N., Nuebler, J., Kanemaki, M.T., Xie, L., Paulson, J.R., Earnshaw, W.C., et al. (2018). A pathway for mitotic chromosome formation. Science 359.

Gullerova, M., and Proudfoot, N.J. (2008). Cohesin complex promotes transcriptional termination between convergent genes in S. pombe. Cell 132, 983–995.

Haarhuis, J.H., Elbatsh, A.M., and Rowland, B.D. (2014). Cohesin and its regulation: on the logic of X-shaped chromosomes. Developmental cell 31, 7–18.

Haarhuis, J.H., Elbatsh, A.M., van den Broek, B., Camps, D., Erkan, H., Jalink, K., Medema, R.H., and Rowland, B.D. (2013). WAPL-mediated removal of cohesin protects against segregation errors and aneuploidy. Current biology: CB 23, 2071–2077.

Haarhuis, J.H.I., van der Weide, R.H., Blomen, V.A., Yanez-Cuna, J.O., Amendola, M., van Ruiten, M.S., Krijger, P.H.L., Teunissen, H., Medema, R.H., van Steensel, B., et al. (2017). The Cohesin Release Factor WAPL Restricts Chromatin Loop Extension. Cell 169, 693–707 e614.

Izumi, K., Nakato, R., Zhang, Z., Edmondson, A.C., Noon, S., Dulik, M.C., Rajagopalan, R., Venditti, C.P., Gripp, K., Samanich, J., et al. (2015). Germline gain-of-function mutations in AFF4 cause a developmental syndrome functionally linking the super elongation complex and cohesin. Nature genetics 47, 338–344.

Jambhekar, A., Emerman, A.B., Schweidenback, C.T., and Blower, M.D. (2014). RNA stimulates Aurora B kinase activity during mitosis. PloS one 9, e100748.

Janicki, S.M., Tsukamoto, T., Salghetti, S.E., Tansey, W.P., Sachidanandam, R., Prasanth, K.V., Ried, T., Shav-Tal, Y., Bertrand, E., Singer, R.H., et al. (2004). From silencing to gene expression: real-time analysis in single cells. Cell 116, 683–698.

Jao, C.Y., and Salic, A. (2008). Exploring RNA transcription and turnover in vivo by using click chemistry. Proceedings of the National Academy of Sciences of the United States of America 105, 15779–15784.

Kagey, M.H., Newman, J.J., Bilodeau, S., Zhan, Y., Orlando, D.A., van Berkum, N.L., Ebmeier, C.C., Goossens, J., Rahl, P.B., Levine, S.S., et al. (2010). Mediator and cohesin connect gene expression and chromatin architecture. Nature 467, 430–435.

Kueng, S., Hegemann, B., Peters, B.H., Lipp, J.J., Schleiffer, A., Mechtler, K., and Peters, J.M. (2006). Wapl controls the dynamic association of cohesin with chromatin. Cell 127, 955–967.

Liang, K., Woodfin, A.R., Slaughter, B.D., Unruh, J.R., Box, A.C., Rickels, R.A., Gao, X., Haug, J.S., Jaspersen, S.L., and Shilatifard, A. (2015). Mitotic Transcriptional Activation: Clearance of Actively Engaged Pol II via Transcriptional Elongation Control in Mitosis. Molecular cell 60, 435–445.

Liu, H., Qu, Q., Warrington, R., Rice, A., Cheng, N., and Yu, H. (2015). Mitotic Transcription Installs Sgo1 at Centromeres to Coordinate Chromosome Segregation. Molecular cell 59, 426–436.

Mannini, L., F, C.L., Cucco, F., Amato, C., Quarantotti, V., Rizzo, I.M., Krantz, I.D., Bilodeau, S., and Musio, A. (2015). Mutant cohesin affects RNA polymerase II regulation in Cornelia de Lange syndrome. Sci Rep 5, 16803.

Miga, K.H., Newton, Y., Jain, M., Altemose, N., Willard, H.F., and Kent, W.J. (2014). Centromere reference models for human chromosomes X and Y satellite arrays. Genome research 24, 697–707.

Molina, O., Vargiu, G., Abad, M.A., Zhiteneva, A., Jeyaprakash, A.A., Masumoto, H., Kouprina, N., Larionov, V., and Earnshaw, W.C. (2016). Epigenetic engineering reveals a balance between histone modifications and transcription in kinetochore maintenance. Nature communications 7, 13334.

Nasmyth, K., and Haering, C.H. (2009). Cohesin: its roles and mechanisms. Annual review of genetics 43, 525–558.

Natsume, T., Kiyomitsu, T., Saga, Y., and Kanemaki, M.T. (2016). Rapid Protein Depletion in Human Cells by Auxin-Inducible Degron Tagging with Short Homology Donors. Cell reports 15, 210–218.

Novais-Cruz, M., Alba Abad, M., van, I.W.F., Galjart, N., Jeyaprakash, A.A., Maiato, H., and Ferras, C. (2018). Mitotic progression, arrest, exit or death relies on centromere structural integrity, rather than de novo transcription. eLife 7.

Olarerin-George, A.O., and Jaffrey, S.R. (2017). MetaPlotR: a Perl/R pipeline for plotting metagenes of nucleotide modifications and other transcriptomic sites. Bioinformatics 33, 1563–1564.

Palazzo, A.F., Springer, M., Shibata, Y., Lee, C.S., Dias, A.P., and Rapoport, T.A. (2007). The signal sequence coding region promotes nuclear export of mRNA. PLoS Biol 5, e322.

Palozola, K.C., Donahue, G., Liu, H., Grant, G.R., Becker, J.S., Cote, A., Yu, H., Raj, A., and Zaret, K.S. (2017). Mitotic transcription and waves of gene reactivation during mitotic exit. Science 358, 119–122.

Prescott, D.M., and Bender, M.A. (1962). Synthesis of RNA and protein during mitosis in mammalian tissue culture cells. Experimental cell research 26, 260–268.

Quinlan, A.R., and Hall, I.M. (2010). BEDTools: a flexible suite of utilities for comparing genomic features. Bioinformatics 26, 841–842.

Rao, S.S.P., Huang, S.C., Glenn St Hilaire, B., Engreitz, J.M., Perez, E.M., Kieffer-Kwon, K.R., Sanborn, A.L., Johnstone, S.E., Bascom, G.D., Bochkov, I.D., et al. (2017). Cohesin Loss Eliminates All Loop Domains. Cell 171, 305–320 e324.

Rosic, S., Kohler, F., and Erhardt, S. (2014). Repetitive centromeric satellite RNA is essential for kinetochore formation and cell division. The Journal of cell biology 207, 335–349.

Schaaf, C.A., Kwak, H., Koenig, A., Misulovin, Z., Gohara, D.W., Watson, A., Zhou, Y., Lis, J.T., and Dorsett, D. (2013). Genome-wide control of RNA polymerase II activity by cohesin. PLoS genetics 9, e1003382.

Segil, N., Guermah, M., Hoffmann, A., Roeder, R.G., and Heintz, N. (1996). Mitotic regulation of TFIID: inhibition of activator-dependent transcription and changes in subcellular localization. Genes & development 10, 2389–2400.

Steurer, B., Janssens, R.C., Geverts, B., Geijer, M.E., Wienholz, F., Theil, A.F., Chang, J., Dealy, S., Pothof, J., van Cappellen, W.A., et al. (2018). Live-cell analysis of endogenous GFP-RPB1 uncovers rapid turnover of initiating and promoter-paused RNA Polymerase II. Proceedings of the National Academy of Sciences of the United States of America 115, E4368–E4376.

Tedeschi, A., Wutz, G., Huet, S., Jaritz, M., Wuensche, A., Schirghuber, E., Davidson, I.F., Tang, W., Cisneros, D.A., Bhaskara, V., et al. (2013). Wapl is an essential regulator of chromatin structure and chromosome segregation. Nature 501, 564–568.

Teves, S.S., An, L., Bhargava-Shah, A., Xie, L., Darzacq, X., and Tjian, R. (2018). A stable mode of bookmarking by TBP recruits RNA polymerase II to mitotic chromosomes. eLife 7.

Titov, D.V., Gilman, B., He, Q.L., Bhat, S., Low, W.K., Dang, Y., Smeaton, M., Demain, A.L., Miller, P.S., Kugel, J.F., et al. (2011). XPB, a subunit of TFIIH, is a target of the natural product triptolide. Nat Chem Biol 7, 182–188.

Trapnell, C., Roberts, A., Goff, L., Pertea, G., Kim, D., Kelley, D.R., Pimentel, H., Salzberg, S.L., Rinn, J.L., and Pachter, L. (2012). Differential gene and transcript expression analysis of RNA-seq experiments with TopHat and Cufflinks. Nature protocols 7, 562–578.

Uhlmann, F., Wernic, D., Poupart, M.A., Koonin, E.V., and Nasmyth, K. (2000). Cleavage of cohesin by the CD clan protease separin triggers anaphase in yeast. Cell 103, 375–386.

Watrin, E., Schleiffer, A., Tanaka, K., Eisenhaber, F., Nasmyth, K., and Peters, J.M. (2006). Human Scc4 is required for cohesin binding to chromatin, sister-chromatid cohesion, and mitotic progression. Current biology: CB 16, 863–874.

Yan, J., Enge, M., Whitington, T., Dave, K., Liu, J., Sur, I., Schmierer, B., Jolma, A., Kivioja, T., Taipale, M., et al. (2013). Transcription factor binding in human cells occurs in dense clusters formed around cohesin anchor sites. Cell 154, 801–813.

