## Supplemental Data for "Cohesin removal reprograms gene expression upon mitotic entry"

### Supplemental Tables

| Name | Specie | Origen | Source | Assay | Dilution |
|---|---|---|---|---|---|
| pS2 | *H. sapiens* | Rabbit | Abcam (5095) | IF/WB/ChIP | 1:400/1:2000 |
| pS5 | *H. sapiens* | Rabbit | Abcam (5131) | IF/WB | 1:400/1:2000 |
| CENPA | *H. sapiens* | Mouse | Abcam (13939) | IF | 1:500 |
| CENPC | *H. sapiens* | Rabbit | Gascoigne et al., 2011 | IF | 1:10000 |
| CENPC | *X. laevis* | Guinea pig | Cocalico (in house, see methods) | IF | 1:500 |
| ACA | *H. sapiens* | Human | Antibodies Inc. (15-234-0001) | IF | 1:500 |
| H3 | *H. sapiens* | Rabbit | Abcam (1791) | WB | 1:5000 |
| GFP | *A. victoria* | Mouse | Abcam (1218) | IF | 1:1000 |
| FLAG | Recombinant | Rabbit | Sigma (F7425) | IF | 1:500 |
| CREST | *H. sapiens* | Human | Inmunovision (HCT-100) | IF | 1:200 |
| RAD21 | *H. sapiens* | Rabbit | Abcam (922) | IF/WB | 1:200/1:2000 |
| SMC3 | *H. sapiens* | Rabbit | Abcam (9263) | IF/WB | 1:400/1:2000 |
| Cherry | DsRed *Discosoma* | Rabbit | Abcam (167453) | WB | 1:2000 |
| WAPL | *X. laevis* | Rabbit | Cocalico (in house, see methods) | WB | 1:1000 |
| WAPL | *H. sapiens* | Rabbit | Abcam (70741) | WB | 1:2000 |
| IgG | | Rabbit | | ChIP | |

**Supplemental Table 1.** List of antibodies used in this study

| **Relative to Figure 1 and Supplemental Figure 2** | | |
|---|---|---|
| **Name** | **Sequence (5'-3')** | **Reference** |
| GAPDH-F | CATGGAGGCCTGGTGGGGGA | This study |
| GAPDH-R | CCGTTGACTCCGACCTTCACC | |
| DXZ1-F | GTGGAGATTTGGACCGCTTTGAG | McNulty et al, 2017 |
| DXZ1-R | CTCAGAGAGGTCCAAATATCCCCT | |
| Alpha3-F | GAGCAGTTTGGAAACCCTCTG | This study |
| Alpha3-R | CTCTGTGAGCCGAATGCACAC | |
| Alpha7-F | AGCGATTTGAGGACAATTGCAG | This study |
| Alpha7-R | GTGAGTTGAACGCACACATCAC | |
| Alpha17-F | GGAAACGGGATAATTTCAGCTGAC | This study |
| Alpha17-R | CACAGAGTGGTCCAAATATCCAC | |
| ACTB-F | GCTATTCTCGCAGCTCACCATGG | This study |
| ACTB-R | ACGATGGAGGGGAAGACGGC | |
| IR-F | GTCCTGCCTGGGGATCAGGTC | This study |
| IR-R | CTGCCTGTTGATGAGGGCTGG | |
| **Relative to Supplemental Figure 9** | | |
| GAPDH-F | CTCATGACCACAGTCCATGCC | This study |
| GAPDH-R | GCCATCCACAGTCTTCTGGGT | |
| MECP2-F | TGAAGGCTGGACACGGAAGCTT | This study |
| MECP2-R | CAGGGATGTGTCGCCTACCTTT | |
| THAP3-F | CGCCTTTGGAAACCGCAAGAAC | This study |
| THAP3-R | CTGAGAAGAGCTGGCATTTCCTC | |
| VPS72-F | GCGAGAAGAGAAGGCACTACTG | This study |
| VPS72-R | CGTTTGTCGTGTATGCTCAGCTG | |
| TET2-F | GCTTACCGAGACGCTGAGGAAA | This study |
| TET2-R | AGAGAAGGAGGCACCACAGGTT | |
| TAF5-F | GCTCCACCTCAGAACAGAATCC | This study |
| TAF5-R | CAAATGGAGGGTAAGCAGTCCG | |
| SF3A2-F | GAAGAACCACCTGGGCTCCTAT | This study |
| SF3A2-R | CAGGTTGGTCTGGTGCTTCTTC | |
| SRRM1-F | CCTCCTCAGAAGATGAACGACC | This study |
| SRRM1-R | GGATGGACTTCTCCTCCGTCTA | |

**Supplemental Table 2.** List of primers used in this study

### Supplemental Figure Legends

**Figure S1. Dynamic of RNA Pol II pS2 and pS5 localization during the interphase-mitosis transition in *Xenopus* egg extracts. A-B**. Chromatin localization of pS2 (A) and pS5 (B) (green) analyzed by immunofluorescence (IF) on DNA spreads from *Xenopus* egg extracts cycled to interphase or subsequently induced for mitotic entry during the indicated times (min). Staining of DNA (blue) and centromere regions (red) was performed using DAPI and anti-CENPC antibodies, respectively. Scale bars, 5 μm. **C-D**. Quantification of pS2 and pS5 centromere fluorescence intensity using images from A-B. Absolute (C) and relative (centromere/total nuclear) levels (D) are plotted. Interphase values were used as reference. In all graphs, error bars indicate the standard deviation (SD) of the mean. *, $P < 0.05$; **, $P < 0.005$; ***, $P < 0.001$ (one-way ANOVA, Dunnett´s test).

**Figure S2. Rad21 knockout eliminates chromosome cohesion and centromeric transcription. A** Analysis of Rad21 localization on metaphase chromosome spreads from HeLa cells expressing sgRNAs targeting the Rad21 gene. Cells were treated with DMSO (control) or doxycycline for 3 days to induce CAS9 expression. Rad21 was detected by IF (red). DAPI staining (blue) and anti-centromere antibodies (CREST; green) were employed for DNA and centromere labeling, respectively. Scale bars, 5 μm. **B** Analysis of Rad21 protein levels by Western blot using total protein extracts isolated from cells treated with DMSO (control) or doxycycline as described above. Levels of histone H3 are provided as loading control. **C** Measurement of cenRNA transcript levels by qPCR in total RNA samples obtained from control (DMSO) or doxycycline-treated cells. The levels of GAPDH were included as a negative control. In all cases, values are normalized to control conditions. Error bars indicate the SD. *, $P < 0.05$; **, $P < 0.005$; ***, $P < 0.001$ (t-test). (ns) no significant differences.

**Figure S3. Selective chromosome arm removal of cohesin during the interphase-mitosis transition in human cells and *Xenopus* egg extracts. A** Smc3 chromatin localization (red) as determined by IF in chromosome spreads from RPE-1 cells in G2, prophase, metaphase or anaphase. Anti-CENPA antibodies (green) and DAPI (blue) were used for centromere and DNA staining, respectively. Scale bars, 10 μm. Measurement of absolute and relative (centromere/total nuclear) Smc3 fluorescence intensity at centromeres is indicated at the bottom. In all cases, values were normalized to G2 levels. Error bars indicate the SD of the mean. *, $P < 0.05$; **, $P < 0.005$; ****, $P < 0.0001$ (one-way ANOVA, Dunnett´s test). **B** Analysis of SMC3 localization (green) by IF using DNA spreads from *Xenopus* egg extracts arrested in interphase and subsequently induced for mitotic entry for the indicated times (min). Anti-CENPC antibodies (red) and DAPI (blue) were used to label centromeres and DNA, respectively. Scale bars, 5 μm. **C-D** Analysis of pS2, pS5 and Smc3 protein levels by Western blotting. Chromatin protein fractions were obtained from human cells (C) or *Xenopus* egg extracts (D) arrested in interphase or induced to enter mitosis for the indicated times (min). Histone H3 levels are provided as loading control.

**Figure S4. Generation and characterization of an AID-WAPL human cell line. A** Schematic representation of the auxin-inducible system used to promote the ubiquitin-mediated degradation of the degron-tagged WAPL protein in HeLa S3 cells. **B** PCR analysis showing the insertion of the AID sequence at the endogenous WAPL locus. **C** Measurement of WAPL levels by Western blotting using total protein extracts from AID-WAPL cells treated with DMSO for 48 h (control) or IAA/Doxycycline for the indicated times. The levels of histone H3 are provided as a loading control. **D** Analysis of metaphase chromosome morphology in AID-WAPL cells treated with DMSO (control) or IAA/Doxycycline for 48 h. Metaphase chromosome spreads were stained with DAPI for DNA labeling. Scale bars, 5 μm. **E** Chromatin localization of Smc3 (red) visualized by IF on metaphase chromosome spreads from AID-WAPL cells treated with DMSO (control) or IAA/Doxycycline for 48 h. Anti-CENPA antibodies (green) and DAPI (blue) were used for centromere and DNA labeling, respectively. Scale bars, 5 μm. **F** Measurement of Smc3 centromere intensity using images from D. Data represent relative enrichment as a centromere/total nuclear ratio and levels were normalized to control (DMSO) conditions. Error bars represent the SD and ***, $P < 0.001$ (t-test). **G** Measurement of Smc3 levels by Western blotting using total or chromatin protein fractions isolated from AID-WAPL cells treated with DMSO (control) or IAA/Doxycycline for 48 h. The levels of histone H3 are included as loading control. **H-I** Analysis of pS5 distribution on chromosome spreads from AID-WAPL cells treated as in E. Anti-CREST antibodies (green) and DAPI (blue) were used for centromere and DNA labeling, respectively. Error bars represent the SD and ns (no significant differences). Scale bars, 5 μm.

**Figure S5. WAPL depletion causes pS2 retention on metaphase chromosomes. A** Analysis of pS2 localization (red) by IF on metaphase chromosome spreads from RPE-1 cells 48 h after transfection with control or WAPL siRNAs. Anti-CENPA antibodies (green) and DAPI (blue) were used for centromere and DNA staining, respectively. Scale bars, 5 μm. **B** Measurement of pS2 centromere signal in images from A. Data represent relative enrichment as centromere/total nuclear signal ratio and levels were normalized to control (non-transfected) conditions. Bars represent the SD and ***, $P < 0.001$ (t-test). (ns) no significant differences. **C-D** Analysis of WAPL, pS2, and Smc3 protein levels by Western blotting using total or chromatin protein fractions from RPE-1 cells transfected with control or WAPL siRNAs for the indicated times. Asynchronous (C) or metaphase-arrested cells (D) were used and in all cases histone H3 levels are provided as loading control. **E** Analysis of WAPL levels by Western blotting using total protein fractions (left) extracted from control (In) and IgG (IgG Δ) or WAPL (WAPL Δ) immuno-depleted egg extracts. WAPL protein levels were also evaluated in IgG and anti-WAPL IP samples to confirm immunoprecipitation of the endogenous protein (right). The levels of histone H3 are shown as loading control. **F-I** Chromatin localization of Smc3 (F) or pS2 (H) (green) by IF on chromosome spreads from control, IgG (IgG Δ), or WAPL (WAPL Δ) immuno-depleted *Xenopus* egg extracts. Anti-CENPC antibodies (red) and DAPI (blue) were used for centromere and DNA staining, respectively. Scale bars, 5 μm. Quantification of Smc3 (G) or pS2 (I) centromere intensity in images from F or H respectively. Data represent relative enrichment as a centromere/total nuclear ratio and levels were normalized to control conditions. Error bars represent the SD and ****, $P < 0.0001$ (t-test). (ns) no significant differences.

**Figure S6. PLK1 and Aurora B inhibition expands chromatin distribution of pS2 in metaphase chromosomes. A** Analysis of pS2 localization (red) by IF using chromosome spreads from RPE-1 cells treated with DMSO (control), barasertib (Aurora B inhibitor) or BI2536 (PLK1 inhibition). DAPI staining (blue) and anti-CENP-A antibodies were used for DNA and centromere labeling, respectively. Scale bars, 5 μm. **B** Quantification of pS2 centromere intensity using images from A. Data indicate relative enrichment as a centromere/total nuclear ratio and levels were normalized to control conditions. **C** Analysis of pS2 localization (green) by IF on chromosome spreads from DMSO-treated (control) *Xenopus* egg extracts or extracts treated with barasertib or BI2536 to inhibit Aurora B or PLK1 kinase activity, respectively. Centromeres (red) and DNA (blue) were labeled using Anti-CENPC antibodies and DAPI staining, respectively. Scale bars, 5 μm. **D** Quantification of pS2 centromere intensity in images from C. Data represent the relative enrichment as centromere/total nuclear ratio and levels were normalized to control conditions. In all graphs, error bars represent the SD of the mean ($n = 3$). **, $P < 0.005$; ***, $P < 0.001$ (t-test).

**Figure S7. Analysis of pS2 localization in metaphase chromosomes of Rad21/WAPL double KO cells. A** pS2 chromatin localization (red) by IF on chromosome spreads from inducible Rad21 mutant HeLa cells transfected with control siRNAs or WAPL siRNAs in the absence of doxycycline (top), treated with doxycycline for 3 consecutive days (Dox), or doxycycline treated cells transfected with siWAPL RNAs (bottom right). Anti-centromere antibodies (CREST; green) and DAPI staining (blue) label centromere and DNA respectively. Scale bars, 10 μm. **B** Quantification of pS2 nuclear intensity using images from A. Levels were normalized to control conditions. Error bars represent the SD of the mean. **, $P < 0.005$; ***, $P < 0.001$; ****, $P < 0.0001$ (t-test).

**Figure S8. Ectopic tethering of the cohesin recruits pS2 and drives chromatin opening at the lacO locus. A-B** Analysis of Smc3 recruitment to the lacO array in U2OS cells. **A** Smc3 chromatin distribution (red) revealed by IF on DNA spreads from U2OS cells transfected with GFP-LacI, Smc1A-LacI or Smc1A*-LacI constructs. Anti-GFP or Anti-FLAG antibodies were used for LacI-tagged protein detection at the lacO locus (green) and DAPI staining for DNA labeling. **B** Quantification of Smc3 intensity at the lacO array in images from A. Measurements were normalized to values obtained following GFP-LacI tethering. **C-D** Analysis of IPTG treatment on the chromatin tethering of LacI-tagged proteins. **C** Analysis of GFP-LacI, Smc1A-LacI or Smc1A*-LacI fusion protein localization (green) revealed by IF in chromosome spreads from U2OS cells transfected with the corresponding constructs in the presence or absence of IPTG. **D** Relative % of cells showing LacI-tagged protein signals at the lacO locus. In all cases, data were normalized to values obtained in the absence of IPTG. **E-H** Analysis of pS2 (E) and pS5 (G) chromatin localization (red) revealed by IF on chromosome spreads from interphase U2OS cells transfected with GFP-LacI, Smc1A-LacI or Smc1A*-LacI constructs. LacI-tagged proteins were detected using Anti-GFP or Anti-FLAG antibodies (green) and DAPI (blue) was used for DNA staining. Quantification of pS2 and pS5 intensity at the lacO array is represented in F and H respectively. **I-J** Analysis of the lacO array size in interphase U2OS cells transfected with GFP-LacI, Smc1A-LacI or Smc1A*-LacI constructs. LacI-tagged proteins and DNA were labeled as described above. In all graphs, error bars represent the SD of the mean (n = 3). *, $P < 0.05$; **, $P < 0.005$; ***, $P < 0.001$ (t-test). In all images, scale bars 10 μm.

**Figure S9. Replicate analysis of gene expression in WAPL-AID cells.**
**A** Metaplot of relative sequencing coverage across all genes expressed at FPKM > 10. **B** Histogram of changes in gene expression between G2 and mitosis in control cells. **C** Heatmap of mitotically repressed genes in control and WAPL-AID cells. **D** Heatmap of mitotically activated genes in control and WAPL-AID cells. **E-G.** Genome browser plots of GMPR2, TAF5, and CRY2 in G2 and mitosis from mock and auxin-treated samples. All genes show mitotic repression in mock but not in auxin-treated samples. Y scale is: -5,20 (GMPR2), -5,20 (TAF5), -5,15 (CRY2).

**Figure S10. Effect of transcriptional inhibitors on metaphase pS2 localization. A** Analysis of pS2 localization (red) in metaphase spreads from AID-WAPL cells incubated with IAA and doxycycline for 48 h, arrested in metaphase, and subsequently treated with triptolide (TRP) for 3 h. Anti-centromere antibodies (CREST; green) and DAPI (blue) were used to stain centromeres and DNA, respectively. Scale bars, 10 μm. **B** Quantification of total nuclear pS2 signal using images from A. Data were normalized to –TRP values. Error bars represent the SD of the mean. **C-E** Analysis of pS2 chromatin localization (red) in metaphase chromosome spreads from RPE-1 cells treated with DMSO (control) or THZ1 or LDC000067 to inhibit Cdk7 or Cdk9 activity, respectively. Anti-CENPA antibodies (green) and DAPI (blue) were used to stain centromeres and DNA, respectively. Scale bars, 5 μm. Quantification of total and centromere pS2 fluorescence intensity using images from C. In all cases, values were normalized to controls and error bars represent the SD of the mean. (ns) no significant differences.

**Figure S11. Analysis of enhancer RNA expression changes in WAPL-AID cells. A** PCA analysis of enhancer expression in mock- and auxin-treated cells showing clearly separated G2 and M cell populations, but no clear separation between mock- and auxin-treated cells. **B-C** Cumulative distribution plots showing the changes in enhancer RNA expression between G2 and mitosis in mock- and auxin-treated cells.

**Figure S12. Analysis of centromere transcription changes in WAPL-AID cells. A** FPKM values for reads aligned to centromere transcript models created using Cufflinks from this sequencing data. FPKM values were compared from mock-treated cells in G2 and mitosis from experiment 1. No significant changes in centromere transcription were detected. **B** Same as A for mock-treated cells in experiment 2. **C** Same as in A for auxin-treated cells from experiment 1. **D** Same as in A for auxin-treated cells for experiment 2. **E** Cumulative distribution plot showing changes in centromere transcripts between G2 and mitosis for mock and auxin-treated cells for experiment 1. **F** Same as in E for experiment 2.

**Figure S13. Modeling GFP-RPB1 fluorescence recovery. A** Fluorescence recovery from control interphase cells plotted with 2- (red) and 3-state (blue) model curves. Residuals after fitting are plotted in lower plot. The 2-state fit does not accurately fit the initial very fast recovery. Modeled parameters and Aikaike Information Criteria and Bayes-Schwarz Information Criteria values for each fit are included as inset. **B** Fluorescence recovery from control interphase cells using normal acquisition parameters. Individuals curve fits and proportion of the recovery explained by each state are indicated next to the plot. Proportions of RNAPII in each state for siControl and siWAPL transfected cells are plotted on the right. **C** Fluorescence recovery was acquired using fast acquisition and fit to a two state model to assess differences in fast recovery dynamics. Recovery and fitted curves as well as measured parameters are indicated on the plot. Proportions of RNAPII in each state for siControl and siWAPL transfected cells are plotted on the right. **D** Fluorescence recovery from siControl transfected cells in mitosis and fitted curves. **E** Fluorescence recovery form siWAPL transfected cells in mitosis and fitted curves.

**Figure S14. Analysis of the contribution of diffusion to GFP-RPB1 fluorescence recovery. A** Analysis of WAPL protein levels by Western blotting in extracts from non-transfected GFP-RPB1 knock-in cells or 48h after transfection with siControl or siWAPL siRNAs. Histone H3 levels are provided as loading control. **B** Snapshot of representative cells after bleaching a 3x3 microns area or half nuclei. The area used for line scanning analysis in the bleached-unbleached transition regions is indicated. Scale bars, 3 $\mu$m. **C-G** Graphs displays average fluorescence intensity versus distance to a reference point (0) at the indicated time points after bleaching ($n \geq 8$). Values were normalized between 0 and 1, RFI (relative fluorescence intensity). Normalized profiles showed a constant shape in interphase cells independently of bleached area size or transfected siRNA, indicating a negligible contribution of diffusion on fluorescence recovery (C-E). A rapid change is observed in the profile shape of mitotic cells transfected with siControl RNAs (F) supporting a major contribution of diffusion. In contrast, profile shape is constant in mitotic cells transfected with siWAPL RNAs (**G**) resembling interphase profiles.

### Methods

#### Cell culture

hTERT RPE-1 (retina epithelium eye) cells were cultured in DMEM:F12 medium supplemented with 10% fetal bovine serum (FBS, Sigma), sodium bicarbonate (0.25%), and 1% penicillin/streptomycin (Corning). HeLa S3 cells were grown in DMEM supplemented with 10% FBS and 1% penicillin/streptomycin. AID-WAPL cell line generation is described below. HeLa-Cas9 cells expressing Rad21 guide RNAs were cultured in DMEM supplemented with tetracycline-free FBS and doxycycline (Sigma D3447, DMSO) added as previously reported to induce Cas9 expression (McKinley and Cheeseman, 2017). MRC-5 fibroblast cells stably expressing GFP-RPB1 were kindly provided by J. Marteijn (Steurer et al., 2018) and cultured in DMEM:F12 with 10% FBS and 1% penicillin/streptomycin. lacO-array-containing U2OS cells is a kind gift from D. Spector and were grown in DMEM supplemented with 10% FBS and 1% penicillin/streptomycin. MDA-MB 435 cells were grown in DMEM supplemented with 10% FBS and 1% penicillin/streptomycin. In all cases cells were incubated at 37° in the presence of 5% $CO_2$.

For G2 arrest RO-3306 (Sigma SML0569, DMSO) was added to a 10μm final concentration for 16h. Cells were washed with 1xPBS and fresh medium added to release cells into mitosis for the indicated times. Metaphase arrest was induced adding nocodazole (Sigma M1404, DMSO) to 0.1μM for 16h. In EU-labelling experiments cells were first synchronized at G1/S using 2mM thymidine (Sigma T1895, water) for 16h, released into fresh medium for 3h and then arrested at G2 or metaphase using RO-3306 or nocodazole. In both cases 5-EU substrate (Abcam ab146642, water) was added to a 0.25mM final concentration for 12h. Aurora-B and Plk1 inhibition was performed adding barasertib or BI-2536 (Selleckchem, DMSO) to a final concentration of 1 and 0.1μm respectively. Inhibition of Cdk7 and Cdk9 kinase activities was induced adding THXZ1 or LDC000067 at 10μm (ApexBio, DMSO) respectively to metaphase arrested cells for 1h. 1μm Triptolide (Sigma T3652, DMSO) was used to inhibit RNAPII transcription initiation in metaphase arrested cells for 3h. In lacO competition assays, IPTG (Sigma I6758, water) was added to a 1mM final concentration for 1h. AID-WAPL proteolytic decay was induced incubating HeLa S3 cells in medium supplemented with 1μg/ml doxycycline (Sigma D3447, DMSO) and 0.5 mM IAA (Indole-3 Acetic Acid sodium salt, Sigma I5148) for 48h.

#### Generation of WAPL-AID cells

To create WAPL-AID cells we first created a repair template that contains a central cassette of mAID-mCherry-P2A-Neomycin (pMB1152). This construct was created using InPhusion cloning from (pMK294, Addgene #72832)*(Natsume et al., 2016)*. We then synthesized 750bp homology regions flanking the WAPL stop codon and fused these to the AID selection cassette using InPhusion cloning. The complete repair template was created by using the InPhusion reaction as a PCR template. PCR product was purified and cotransfected into HeLa S3 cells with a plasmid expressing a WAPL sgRNA and Cas9 (cloned into pX330, Addgene# 42230) (pMB1155). Cells were allowed to grow for 3 days then were split into 500ug/ml Neomycin for ~14 days. Single cell colonies were isolated using cloning rings. Single colonies were initially screened for nuclear mCherry fluorescence, then by Western blot with antibodies against *Xenopus* WAPL that cross react with human WAPL. Potential homozygous knock-in cell lines were then screened by PCR with primers that recognize both the endogenous stop codon or primers that recognize the knock-in junction.

**siRNA and DNA transfection**

ON-TARGETplus Human WAPL siRNA-SMARTpool (Dharmacon, 23063) were transfected into RPE-1 or HeLa cells using the RNAiMAX transfection reagent (Thermofisher 13778030). 20μM siRNAs aliquots were incubated with transfection agent in Optimem medium at room temperature for 20 minutes and added to cells grown in medium without antibiotics. siGENOME non-targeting siRNA (Dharmacon, D-001210) was transfected in parallel as a negative control. The effects of siRNAs were evaluated 48 hours after transfection.

Plasmid DNA transfection was performed in all cases using the Lipofectamine 3000 reagent (Thermofisher, L3000001). DNA (2.5-5 μg) and transfection reagent were incubated in Optimem medium at room temperature for 20 minutes and then added to the cell medium. The expression of transfected DNA was analyzed 48-72 hours after transfection.

**Egg extracts from *Xenopus laevis***

CSF-arrested *Xenopus* egg extracts were prepared and utilized for immunofluorescence as described previously (Hannak and Heald, 2006). Sperm nuclei were cycled through interphase by the addition of 0.6 mM $CaCl_2$ for 90 minutes, then driven back into mitosis by adding an equal volume of CSF-arrested extract for the indicated times. AuroraB and Plk1 inhibition was performed adding barasertib or BI-2536 to a final concentration of 10 and 1μm respectively.

WAPL inmunodepletion was performed using a customized in-house antibody (see below) coupled to protein A dynabeads (Life technologies) and added at 0.125μm/μl egg extract. IgG-coupled beads were used at same concentration in parallel as control. Reactions were incubated at 4° for 1h before being used for immunofluorescence or western blot.

**Recombinant protein and Antibody generation**

A N-terminal fragment corresponding to 1-466 amino acids (~ 46 KDa) of *Xenopus* WAPL protein was cloned to express a GFP-WAPL-6xHis fusion protein. For CENPC expression a fragment containing 381-712 amino acids (~ 32 KDa) of *Xenopus* CENPC protein was used. In both cases, expression was performed in BL21 Rosetta cells as previously described *(5)* and purified using NiNTA and Superdex S200. Peak fractions from S200 were incubated with Precision Protease (GE Healthcare, 27084301) overnight at 4° to excise GFP. Cleaved WAPL/CENPC was bound to HiTrap Heparin and eluted with a linear gradient of NaCl (from 150 mM to 1 M). Fractions containing WAPL/CENPC were loaded into acrylamide gel and a 46/33 KDa band excised and used for antibody generation in rabbit or guinea pig respectively (Cocalico Biologicals). Antibodies were purified from serum by affinity purification using the same recombinant proteins employed for animal injection and the concentration estimated measuring absorbance at 280nm. Specificity was tested by WB using total protein samples from

*Xenopus* egg extracts revealing bands corresponding to the full length of WAPL or CENPC proteins.

**Immunofluorescence**

A list of antibodies used in this study is displayed in Supplemental table1. For chromosome spreads 500μl of cells ($2x10^5$cells/ml) were centrifuged in a cytospin (1800rpm, 10min). Slides were fixed in 1xPBS containing 4% PFA for 10min followed by permeabilization in 1xPBS with 0.5% Triton X-100 for 20min and washed twice with 1xPBS. Slides were blocked in 1xPBS supplemented with 10% ultrapure BSA (Invitrogen, AM2616) for 10min before primary and secondary antibody incubations at 37°. For pS2 and pS5 immunostaining an unfixed KCM protocol was used instead (Chan et al., 2012). Cells ($2x10^5$cells/ml) were swelled in KCl (65mM) at 37° for 10min before cytospin. Slides were incubated in KCM buffer supplemented with 10% ultrapure BSA for 10min prior to primary and secondary antibody incubations performed at 4° for 1h. Slides were finally washed with KCM, fixed in 4% PFA KCM and mounted with DAPI containing solution (Sigma, DUO82040). For EU-labelled RNA visualization the Click-iT RNA Imaging Kit was used following the manual specifications (Invitrogen, C10329). Alexa Fluor 488 azide was employed in the Click-iT reaction performed with fixed cells previously incubated with or without 0.25mM EU.

*Xenopus* egg extracts were fixed in 1xPBS with 4% PFA for 10min prior to centrifugation (10200rpm, 20min) through a cushion buffer containing 1xBrB, 30% glycerol and 0.1% Triton-X-100. Coverslips were washed once in 1xPBS prior to blocking in 1xPBS containing 0.2% Tween and 10% BSA. Incubation of primary antibodies were performed at 4° overnight and secondary at room temperature during 1h. DAPI containing medium was used for mounting.

In all cases samples were storage at 4° under dark conditions until image acquisition.

**Image acquisition, analysis and plotting**

All images were acquired using the microscope set-up described previously (Jambhekar et al., 2014). In all cases images were captured from at least three independent experiments considering 10-20 cells per replicate. This numbers were significantly increased in the experiments with the double RAD21/WAPL mutant cells. For quantitative analysis all images were captured using identical conditions and are displayed following identical settings. Previous to the analysis all images were background-subtracted using Image J software. To delimit inner centromere CENPA, CENPC and ACA antibodies were employed to stain kinetochores and a 2.5μm diameter circle including both sister kinetochores was drawn. Total chromatin signal was measured using DAPI staining as reference and relative centromere measurement were calculated normalizing average centromere to average total nuclear signal. In lacO-tethering experiments an equal size circle was drawn to delimit the lacO area detected by tethering GFP-, SMC1A- or SMC1A*-LacI fusion proteins. DAPI signal was used as reference to measure total chromatin levels and to normalize and calculate average relative signal values at lacO.

Statistical analyses and plotting of image measurements were performed using Prism 7 software. Differences between two groups of data were analyzed by a single-sample t-test. For multiple group analyses one-way ANOVA was performed and Dunnett's comparisons test considered.

**Fluorescence recovery after photobleaching (FRAP)**

All images were acquired using a Ti-2 Eclipse microscope (Nikon) and a previously described GFP-RPB1 knock-in cell line (Steurer et al., 2018) 48h after transfection with siControl or siWapl siRNAs as detailed above. All images were acquired at 37° using a temperature-controlled Tokai-Hit culture dish system. A square of 3x3 microns was imaged during 2 s at 1 s/frame before stimulation for photobleaching using a 488-nm laser at 40% power during 2 s. Time-lapse imaging was performed in two phases: a fast acquisition at 500 ms/frame during 30 s following by a 1 s/frame phase for 3 min and 30 s. For fast recovery analysis images were taken at 300 ms/frame during 30 s followed by a 1 s/frame phase of 30 s. Images were background subtracted and fluorescence intensity evaluated at bleached areas using Fiji software. Measurements were normalized to values obtained before bleaching (t = 0) and related to an unbleached area used as a reference. All recovery graphs show relative fluorescence intensity (RFI) in time (s). In diffusion-dependence tests half nuclei or squares of 3x3 microns were stimulated and imaged as described above. Images were background subtracted and fluorescence intensity evaluated at the indicated time frames in the bleached/unbleached transition zone by line scanning as described previously (Beaudouin et al., 2006; Mueller et al., 2008). RFI values were plotted versus the distance (microns) to an arbitrary point set at the edge of the bleached area.

Normalized FRAP data were fit to two or three component, reaction-dominant models for recovery (Sprague et al., 2004) using the non-linear least-squares method in R. We determined the appropriate number of components for each model by examining the distribution of residuals for each fit and by calculating the Akaike Information Criteria (AIC) and Bayes-Schwarz Information Criteria (BIC) for each model as described (Darzacq et al., 2007).

**Western Blotting**

Total protein from human cells was extracted incubating cells in lysis buffer (150 mM KCl, 25 mM Tris pH 7.4, 5 mM EDTA, 5 mM $MgCl_2$, 1% NP 40, 0.5 mM DTT, 1 mM PMSF) supplemented with protease inhibitors (Roche). Protein concentrations were calculated by Bradford and 10 μg were typically loaded for PAGE-SDS. Proteins were transferred to nitrocellulose membranes (Amersham Protran 0.45 μm NC) using Tris/Glycine buffer with 20% methanol for 1h at room temperature or overnight at 4°. Membranes were then blocked in 1x TBST supplemented with 5% skim milk powder (Millipore). Antibodies were diluted in TBST 5% milk and incubated at 4° overnight (primary) or 1 hour (secondary). Membranes were finally developed using Western Lighting Plus ECL (Perkin Elmer) and scanned in a Chemidoc MP imager (BioRad). Total protein from *Xenopus* extracts were obtained following standard protocols and employed in western blot as described above.

Chromatin-associated protein fractions from human cells or *Xenopus* egg extracts were purified following previously described protocols (MacCallum et al., 2002; Mendez and Stillman, 2000).

**RNA purification**

Total RNA from human cells (~ $10^6$ cells) was extracted using Trizol (Invitrogen, 15596026). Cells were collected and washed in 1xPBS prior to resuspension in 1ml Trizol. 250μl of chloroform was then added for protein extraction following RNA precipitation with isopropanol. RNA pellet was finally washed in 70% ethanol and resuspended in nuclease-free water (Ambion, AM9932). RNA levels were measured using a Biodrop and storage at -20°.

EU-labelled RNA was purified from total RNA extracted from G2 and metaphase arrested cells incubated with 0.25mM EU for 12 hours. For EU-RNAseq we previously removed the rRNA from 5μg total RNA using Ribo-Zero rRNA Removal Kit (Illumina) following the manual specifications. EU-labelled RNA was then recovered from depleted RNA (~ 500 ng) using the Click-iT RNA Capture Kit (Invitrogen, C10365). Renilla and Luciferase custom biotinylated probes were used as spike in controls in the EU-RNAseq experiment. Templates DNA were generated by PCR and *in-vitro* transcribed using the Hiscribe T7 kit (NEB E2030) including biotinylated UTP. Probes were purified by LiCl precipitation and added after the Click-iT reaction at 0.25ng/μl and 0.025 ng/μl final concentration respectively. Purified EU-RNA was used for either cDNA synthesis or library preparation (see above).

**Library preparation**

EU-RNA from G2 and metaphase arrested cells treated or not with Auxin and doxycycline were analyzed in duplicated (Auxin vs Mock). For library construct we used EU-labelled RNA purified from ribo-depleted samples as described above and the NEBNext Ultra Directional RNA Library Prep Kit (NEB) following the manual specifications. Amplified libraries sizes were evaluated by loading into an 8% native PAGE and gel bands excised and purified overnight in a $NH_4Ac$/SDS (0.45 M/0.045%) solution. Libraries were finally precipitated with isopropanol and Glycogen (Thermofisher, 10814010), dissolved in nuclease-free water and submitted to the MGH-sequencing facility for QC and Illumina sequencing.

**EU-RNAseq analysis**

Fastq files from the Illumina HiSeq were collapsed to unique reads using a custom Perl script. Reads were aligned to the human genome (hg38) or human centromere sequences *(Miga et al., 2014)* (released with hg38) using TopHat2. Reads mapping the Gencode gene models were counted using the cuffdiff function in Cufflinks *(Trapnell et al., 2012)*. For analysis of centromere transcripts, we used all reads (from 8 libraries) - mapped to the centromere models as input for Cufflinks to build transcripts. We then counted mapping reads against the Cufflinks gene models. To calculate the reads mapping to enhancers we downloaded the GeneHancer *(Fishilevich et al., 2017)* models from UCSC and removed all enhancers contained within genes using the bedtools *(Quinlan and Hall, 2010)* subtract command. Reads were then counted against intergenic enhancers using cuffdiff. All calculations were performed on FPKM normalized values. For analysis of genes we analyzed all genes with a FPKM value > 10 in all G2 samples and > 0 in all mitosis samples. For analysis of enhancers we analyzed all enhancers with a FPKM value > 1 in all G2 samples and > 0 in all mitosis samples. For analysis of centromere transcripts, we did not impose a FPKM cutoff prior

to analysis. Genome browser images were produced in R using the Gviz package. All other plots and statistical tests were prepared in R.
All sequence files were deposited into GEO (https://www.ncbi.nlm.nih.gov/geo/query/acc.cgi?acc=GSE130558).

We attempted to use Renilla and Firefly spike-in RNAs to normalize our EU-RNA-Seq data. However, in one sample the Firefly and Renilla spike-in sequences diverged dramatically and were not usable, suggesting a gene specific effect as observed for spike-in sequences in previous studies (Consortium, 2014; Risso et al., 2014). Additionally, we performed Q-RT-PCR validation experiments of additional EU-labeled biological replicates. In all cases validation analysis corresponded to expression values/ratios obtained using bulk FPKM normalization and not to normalization obtained by spike-in normalization. As a result all our library comparisons were performed with FPKM normalized samples.

To produce metaplots of sequencing coverage across genes we adapted the metaPlotR software *(Olarerin-George and Jaffrey, 2017)*. We created gene models for each human gene expressed at FPKM > 10, including introns, using R. We then converted our mapped reads into bedgraph files using BEDTools. These files were used as input for metaPlotR software. Metaplots were produced using ggplot in R.

**RT-qPCR**

Total or EU-labelled RNA (~ 500 ng) was reverse transcribed using the iScript cDNA synthesis kit (Bio-Rad, 1708890) following the protocol specifications. cDNA was then diluted and used as template in 15μl final volume RT reactions containing IQ-Sybr Green reagent (Bio-Rad, 1708880). Water and -RT samples were included as negative controls. A list of primers used in this experiments is displayed in supplemental table 2.

**ChIP-qPCR**

Chromatin immunoprecipitation assay was performed essentially following the protocol reported by Myers (v011014)( https://hudsonalpha.org/protocols/). Briefly, G2 and metaphase arrested RPE-1 cells were collected and fixed in 1xPBS containing 1% formaldehyde for 10 minutes. Glycine was added to a final concentration of 0.125M to stop cross-linking and cells washed in cold 1xPBS. Intact nuclei from ~ $5x10^6$ cells were extracted with 1ml of Farnham lysis buffer and finally resuspended into 300μl of RIPA buffer. Nuclei were fragmented using a Qsonica sonicator for 5 minutes (30 seconds on/off, 40%). The efficiency of fragmentation was measured as the enrichment of ~ 300 bp fragments, tested on an agarose gel, and the DNA quantified in a Biodrop. To perform immunoprecipitations Dynabeads M-280 Sheep Anti-Rabbit (Invitrogen 11203D) were coupled to 5μg of pS2 or IgG rabbit antibodies and incubated with 25μg of fragmented chromatin samples overnight at 4°. Beads were then washed with LiCl wash buffer and eluted in 200μl of IP elution buffer. Cross-link was reversed by incubating beads at 65° for 1 hour, supernatant (DNA) recovered after centrifugation and incubated at 65° for 14h. In parallel, input samples (25μg of fragmented chromatin) were incubated at 65° for 14h and then purified using QIAquick PCR Purification Kit (QIAGEN). Serial dilutions of Input and recovered DNA samples were used as template in 15μl RT reactions containing iQ-Sybr Green reagent (Bio-Rad). In all the assays water was included as negative control and data represented as the percentage of input recovered in each condition. The analysis of an intergenic region (IR) was included as a negative control. The primers employed in this assay are listed in supplemental table 2.

**Plasmids and constructs**

For expression of SMC1A-LacI fusion proteins we used the pGK215 vector (Addgene 45110) (Gascoigne et al., 2011) where GFP encoding DNA was replaced by the full-length cDNA encoding the SMC1A protein (pMB1208). A point mutation was then introduced to express a modified SMC1A version (SMC1A*) containing an amino acid substitution (L1128V)(pMB1209) that generates a WAPL-resistant cohesin ring (Elbatsh et al., 2016). In both cases a C-terminal FLAG was added for further detection. The SV40 promoter was finally cloned to drive the expression of GFP-, SMC1A- or SMC1A*-LacI fusion proteins in mammalian cells. All DNA amplifications were performed using the Q5 High Fidelity polymerase (NEB). For constructs related to the AID-WAPL cell line see above.

For protein expression in bacteria DNA fragments encoding WAPL (1-466aa) and CENPC (381-712aa) from *Xenopus laevis* were cloned into the pET30a-GFP plasmid to express GFP/6xHis-tagged fusion proteins.

### References


Beaudouin, J., Mora-Bermudez, F., Klee, T., Daigle, N., and Ellenberg, J. (2006). Dissecting the contribution of diffusion and interactions to the mobility of nuclear proteins. Biophys J *90*, 1878-1894.

Chan, F.L., Marshall, O.J., Saffery, R., Kim, B.W., Earle, E., Choo, K.H., and Wong, L.H. (2012). Active transcription and essential role of RNA polymerase II at the centromere during mitosis. Proceedings of the National Academy of Sciences of the United States of America *109*, 1979-1984.

Consortium, S.M.-I. (2014). A comprehensive assessment of RNA-seq accuracy, reproducibility and information content by the Sequencing Quality Control Consortium. Nat Biotechnol *32*, 903-914.

Darzacq, X., Shav-Tal, Y., de Turris, V., Brody, Y., Shenoy, S.M., Phair, R.D., and Singer, R.H. (2007). In vivo dynamics of RNA polymerase II transcription. Nature structural & molecular biology *14*, 796-806.

Elbatsh, A.M.O., Haarhuis, J.H.I., Petela, N., Chapard, C., Fish, A., Celie, P.H., Stadnik, M., Ristic, D., Wyman, C., Medema, R.H.*, et al.* (2016). Cohesin Releases DNA through Asymmetric ATPase-Driven Ring Opening. Molecular cell *61*, 575-588.

Fishilevich, S., Nudel, R., Rappaport, N., Hadar, R., Plaschkes, I., Iny Stein, T., Rosen, N., Kohn, A., Twik, M., Safran, M.*, et al.* (2017). GeneHancer: genome-wide integration of enhancers and target genes in GeneCards. Database (Oxford) *2017*.

Gascoigne, K.E., Takeuchi, K., Suzuki, A., Hori, T., Fukagawa, T., and Cheeseman, I.M. (2011). Induced ectopic kinetochore assembly bypasses the requirement for CENP-A nucleosomes. Cell *145*, 410-422.

Hannak, E., and Heald, R. (2006). Investigating mitotic spindle assembly and function in vitro using Xenopus laevis egg extracts. Nature protocols *1*, 2305-2314.

Jambhekar, A., Emerman, A.B., Schweidenback, C.T., and Blower, M.D. (2014). RNA stimulates Aurora B kinase activity during mitosis. PloS one *9*, e100748.

MacCallum, D.E., Losada, A., Kobayashi, R., and Hirano, T. (2002). ISWI remodeling complexes in Xenopus egg extracts: identification as major chromosomal components that are regulated by INCENP-aurora B. Molecular biology of the cell *13*, 25-39.
McKinley, K.L., and Cheeseman, I.M. (2017). Large-Scale Analysis of CRISPR/Cas9 Cell-Cycle Knockouts Reveals the Diversity of p53-Dependent Responses to Cell-Cycle Defects. Developmental cell *40*, 405-420 e402.
Mendez, J., and Stillman, B. (2000). Chromatin association of human origin recognition complex, cdc6, and minichromosome maintenance proteins during the cell cycle: assembly of prereplication complexes in late mitosis. Molecular and cellular biology *20*, 8602-8612.
Miga, K.H., Newton, Y., Jain, M., Altemose, N., Willard, H.F., and Kent, W.J. (2014). Centromere reference models for human chromosomes X and Y satellite arrays. Genome research *24*, 697-707.
Mueller, F., Wach, P., and McNally, J.G. (2008). Evidence for a common mode of transcription factor interaction with chromatin as revealed by improved quantitative fluorescence recovery after photobleaching. Biophys J *94*, 3323-3339.
Natsume, T., Kiyomitsu, T., Saga, Y., and Kanemaki, M.T. (2016). Rapid Protein Depletion in Human Cells by Auxin-Inducible Degron Tagging with Short Homology Donors. Cell reports *15*, 210-218.
Olarerin-George, A.O., and Jaffrey, S.R. (2017). MetaPlotR: a Perl/R pipeline for plotting metagenes of nucleotide modifications and other transcriptomic sites. Bioinformatics *33*, 1563-1564.
Quinlan, A.R., and Hall, I.M. (2010). BEDTools: a flexible suite of utilities for comparing genomic features. Bioinformatics *26*, 841-842.
Risso, D., Ngai, J., Speed, T.P., and Dudoit, S. (2014). Normalization of RNA-seq data using factor analysis of control genes or samples. Nat Biotechnol *32*, 896-902.
Sprague, B.L., Pego, R.L., Stavreva, D.A., and McNally, J.G. (2004). Analysis of binding reactions by fluorescence recovery after photobleaching. Biophys J *86*, 3473-3495.
Steurer, B., Janssens, R.C., Geverts, B., Geijer, M.E., Wienholz, F., Theil, A.F., Chang, J., Dealy, S., Pothof, J., van Cappellen, W.A.*, et al.* (2018). Live-cell analysis of endogenous GFP-RPB1 uncovers rapid turnover of initiating and promoter-paused RNA Polymerase II. Proceedings of the National Academy of Sciences of the United States of America *115*, E4368-E4376.
Trapnell, C., Roberts, A., Goff, L., Pertea, G., Kim, D., Kelley, D.R., Pimentel, H., Salzberg, S.L., Rinn, J.L., and Pachter, L. (2012). Differential gene and transcript expression analysis of RNA-seq experiments with TopHat and Cufflinks. Nature protocols *7*, 562-578.

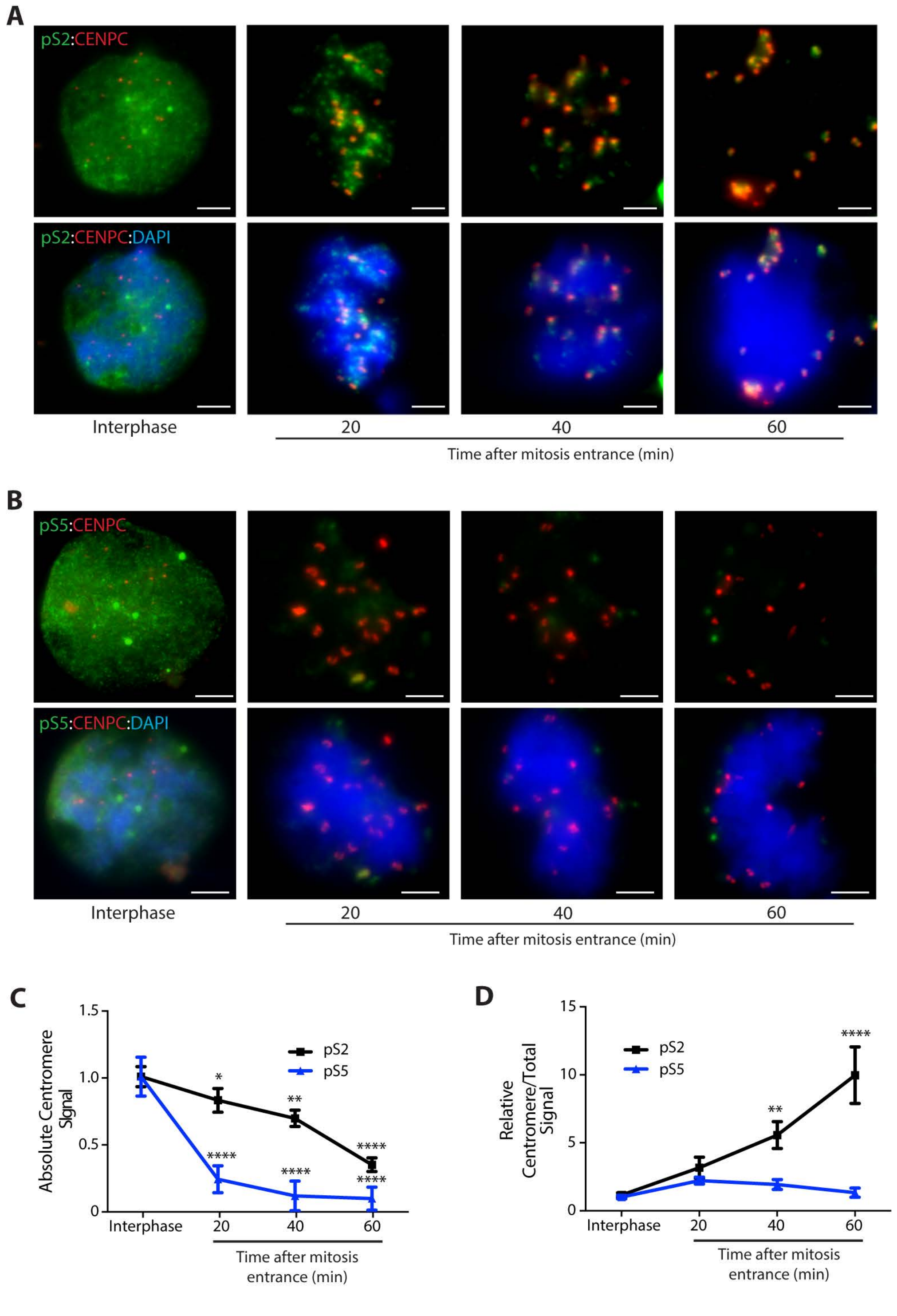


**Supplemental Figure 1**

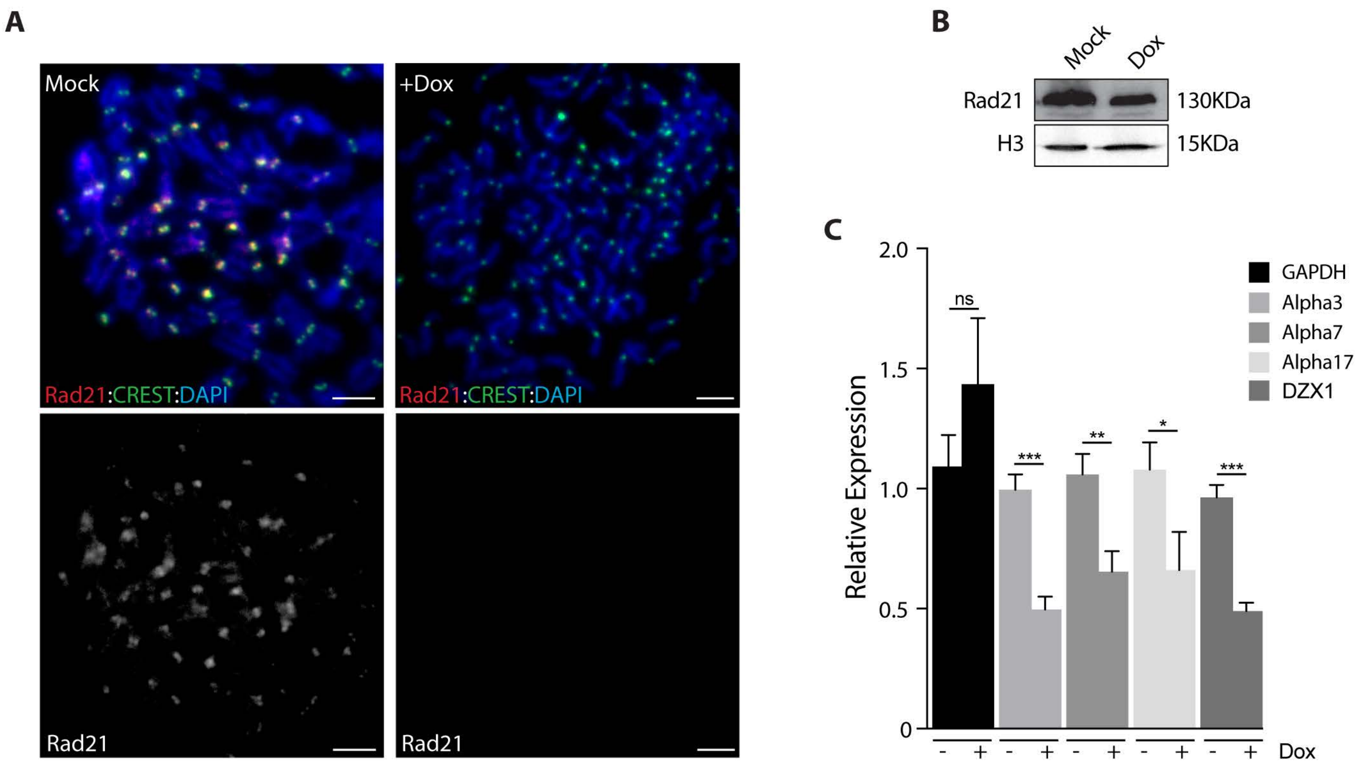


**Supplemental Figure 2**

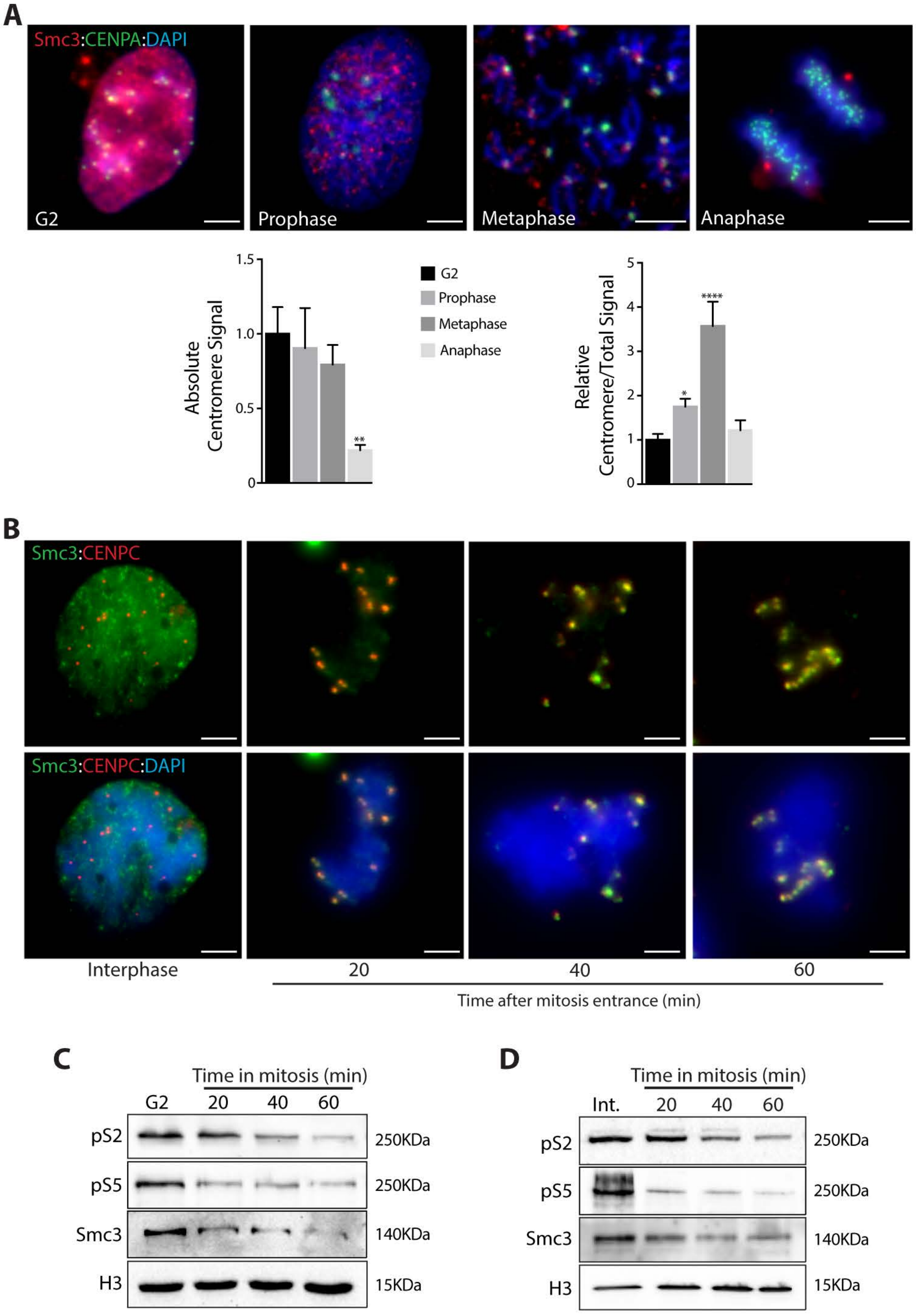


**Supplemental Figure 3**

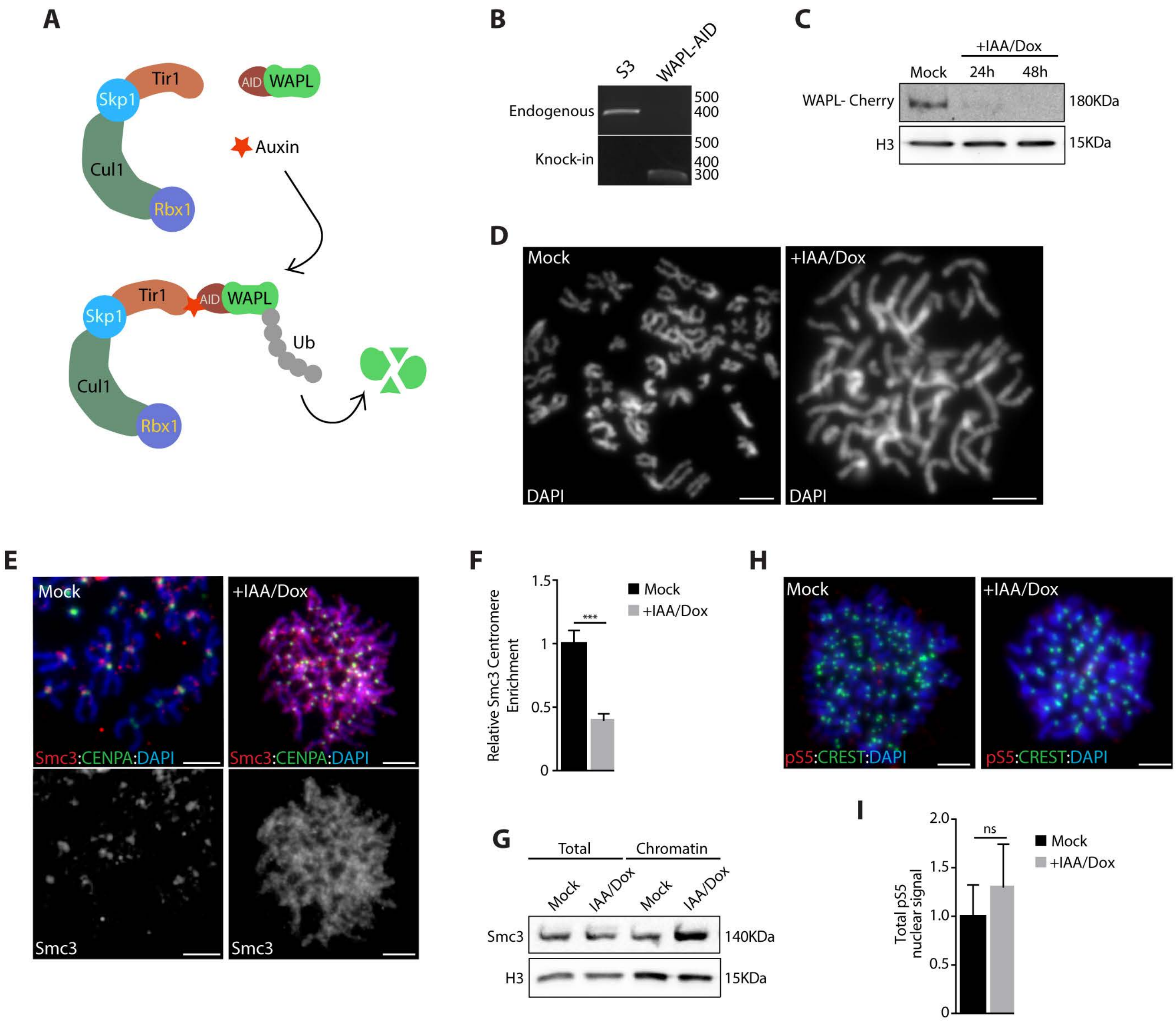


**Supplemental Figure 4**

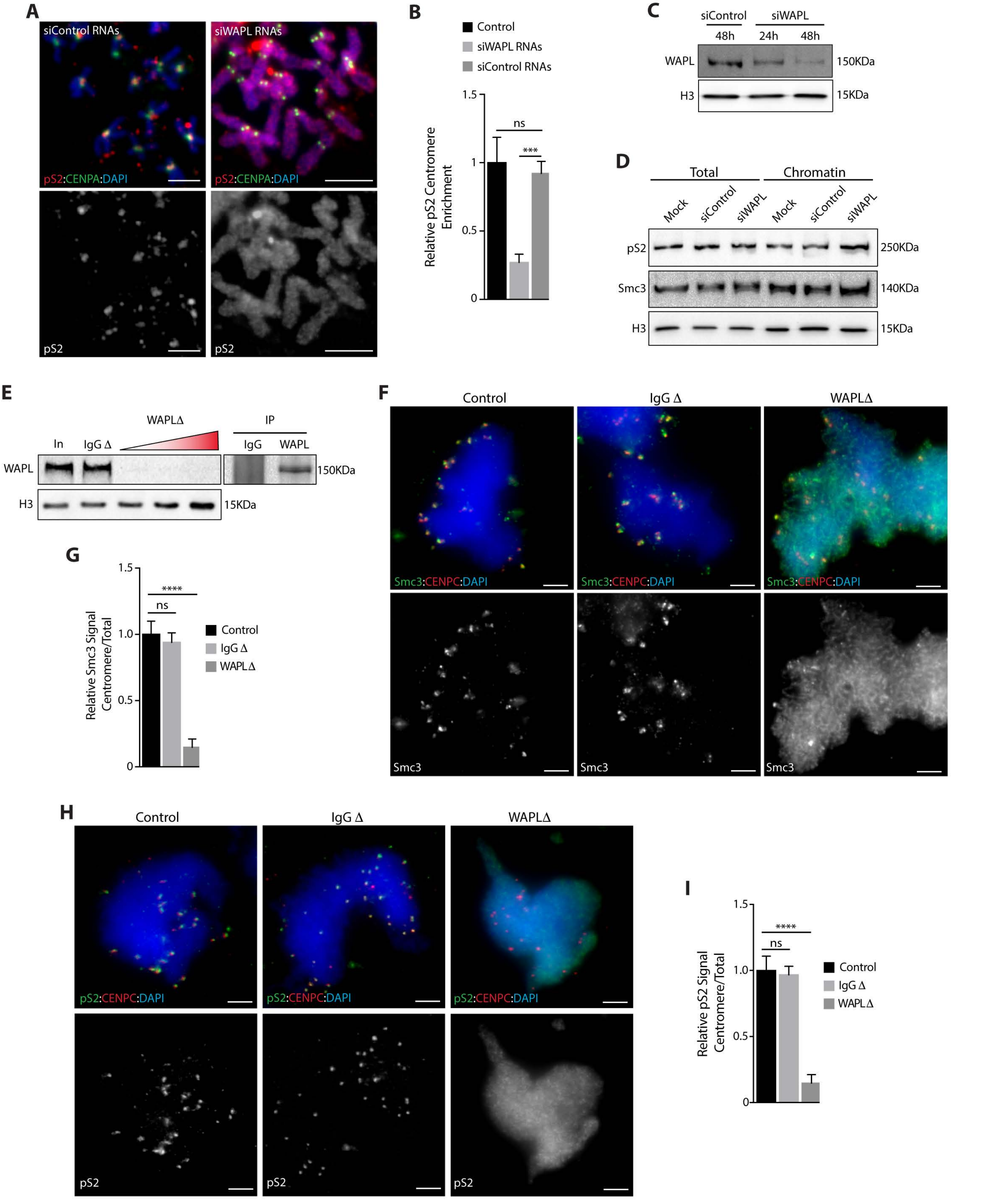


**Supplemental Figure 5**

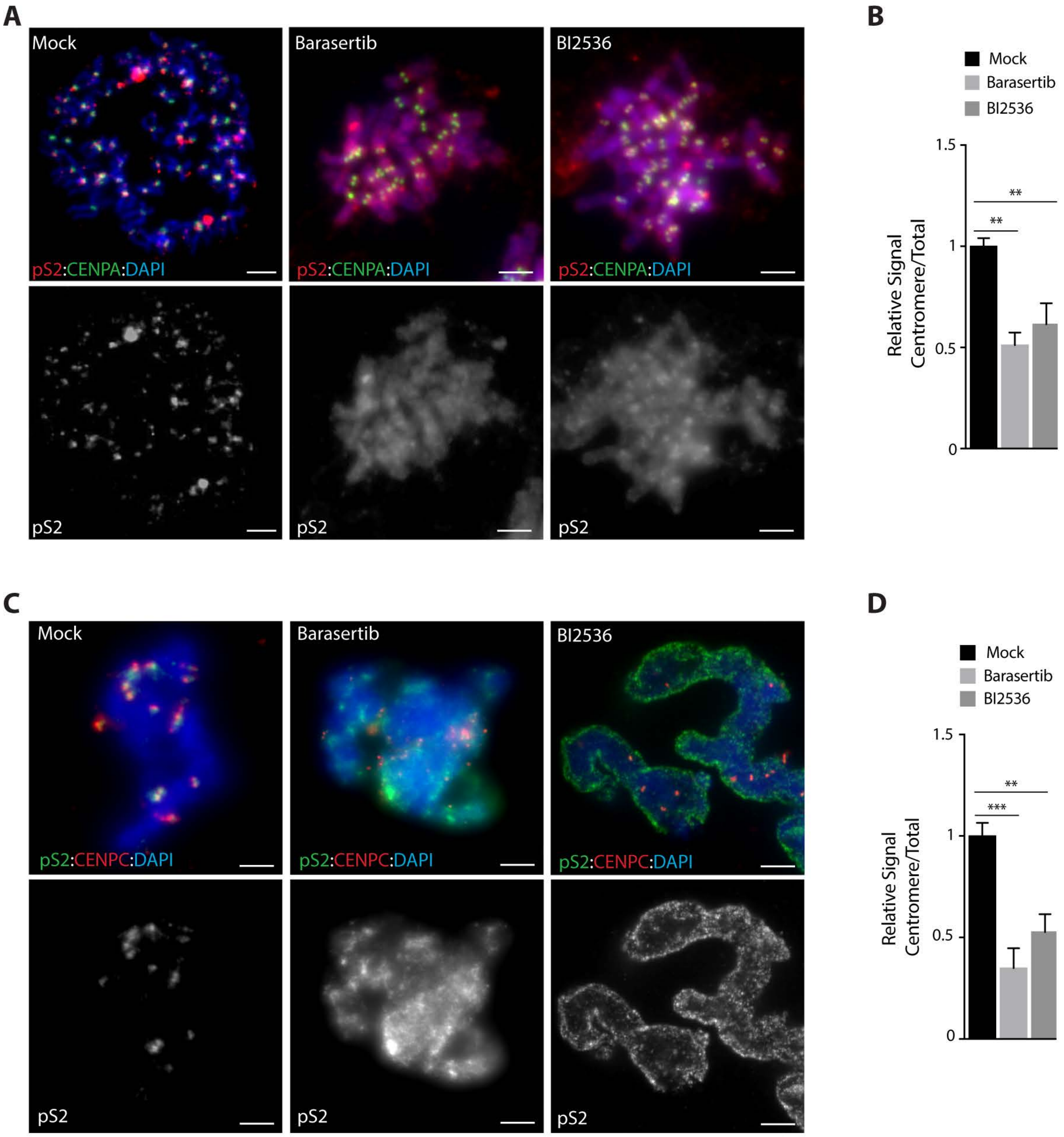


**Supplemental Figure 6**

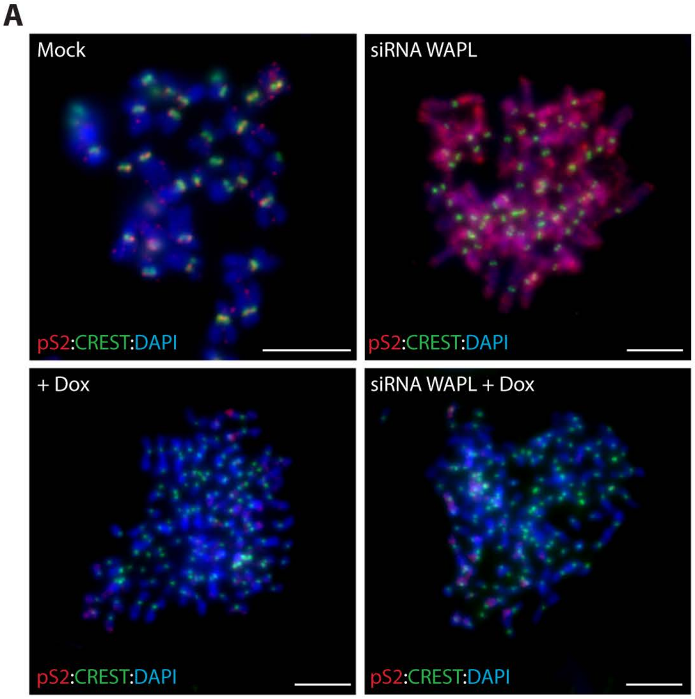


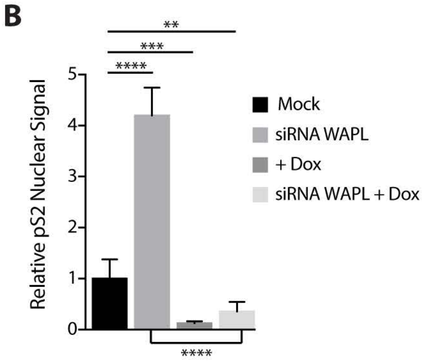


**Supplemental Figure 7**

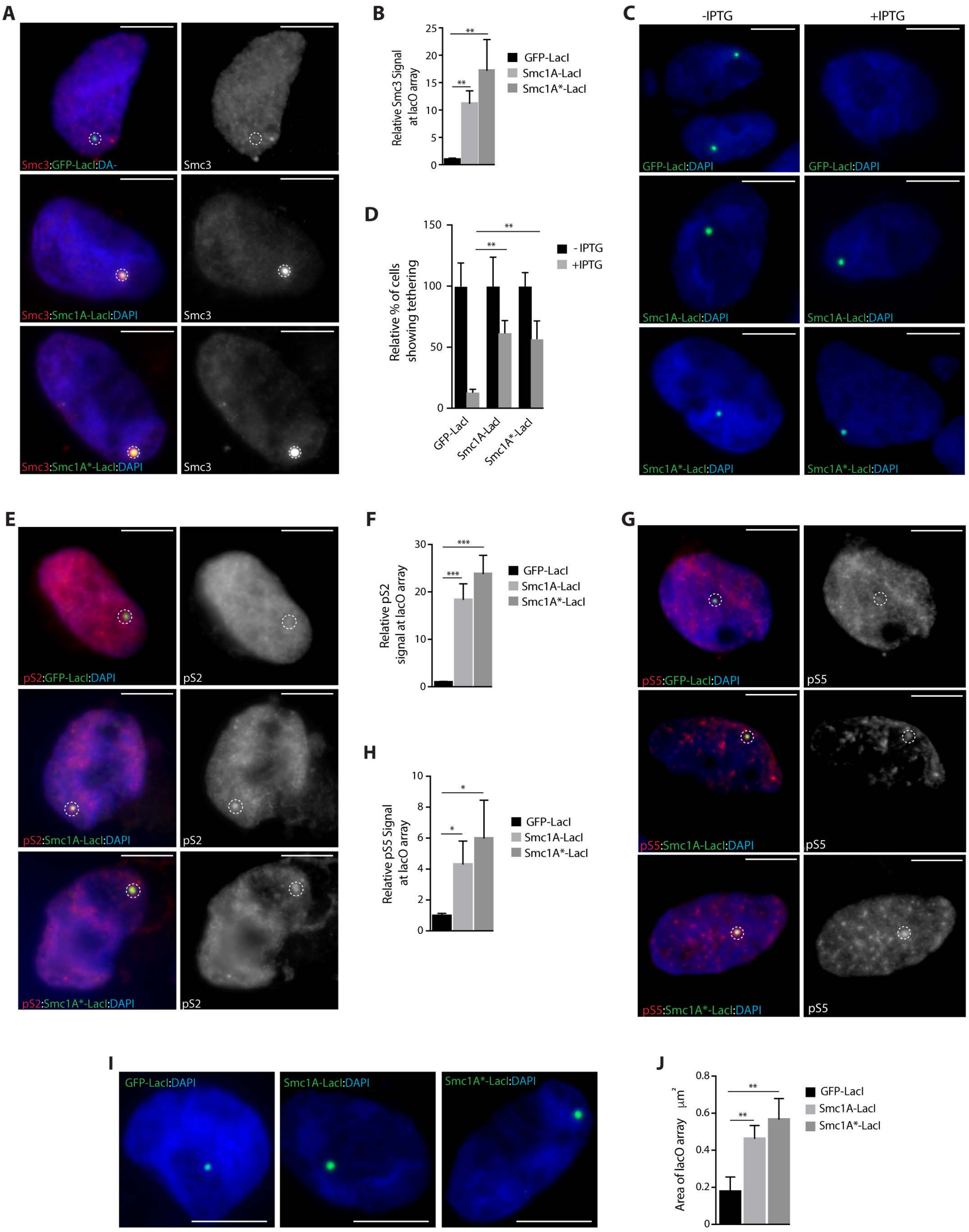


**Supplemental Figure 8**

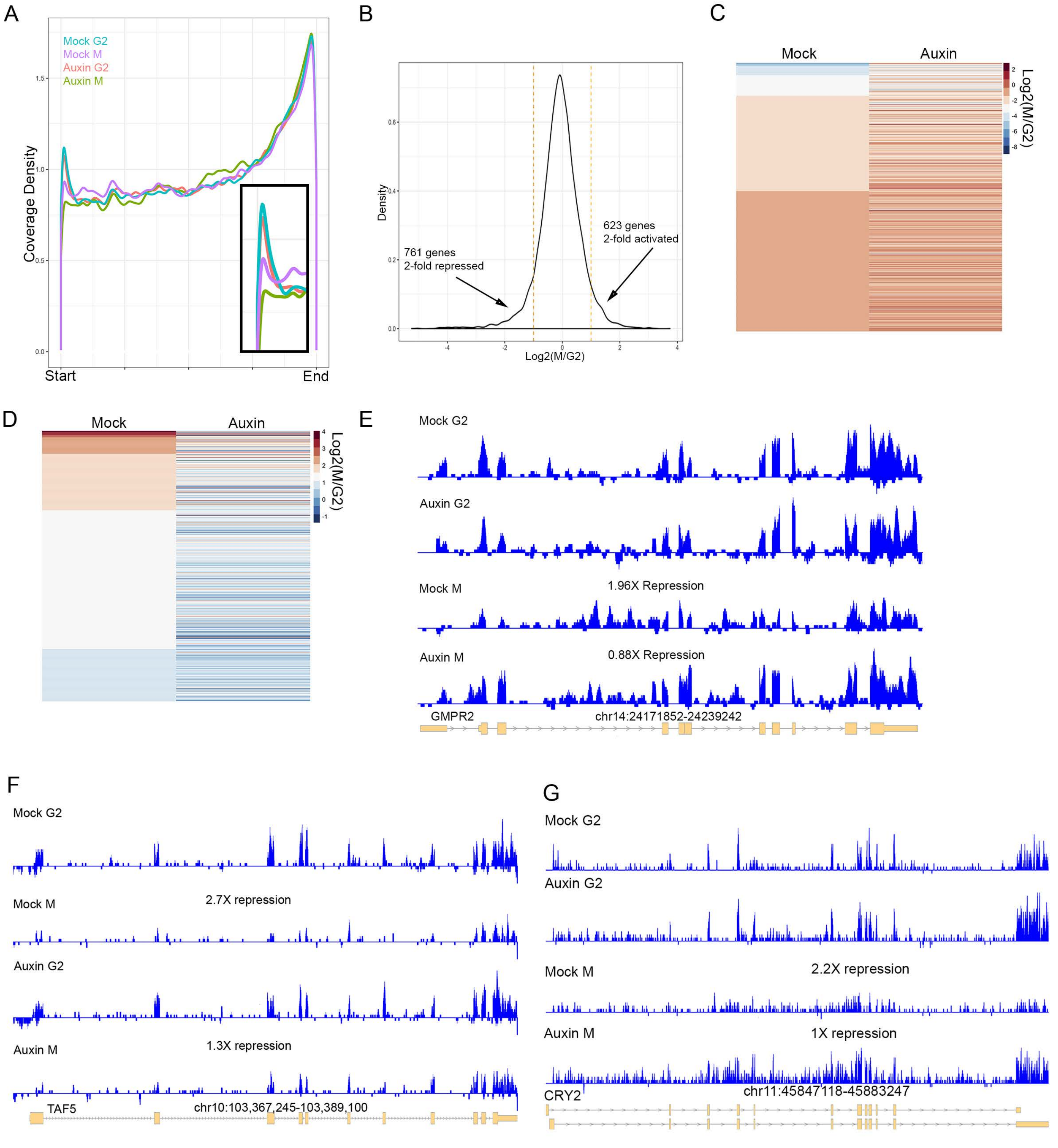


**Supplemental Figure 9**

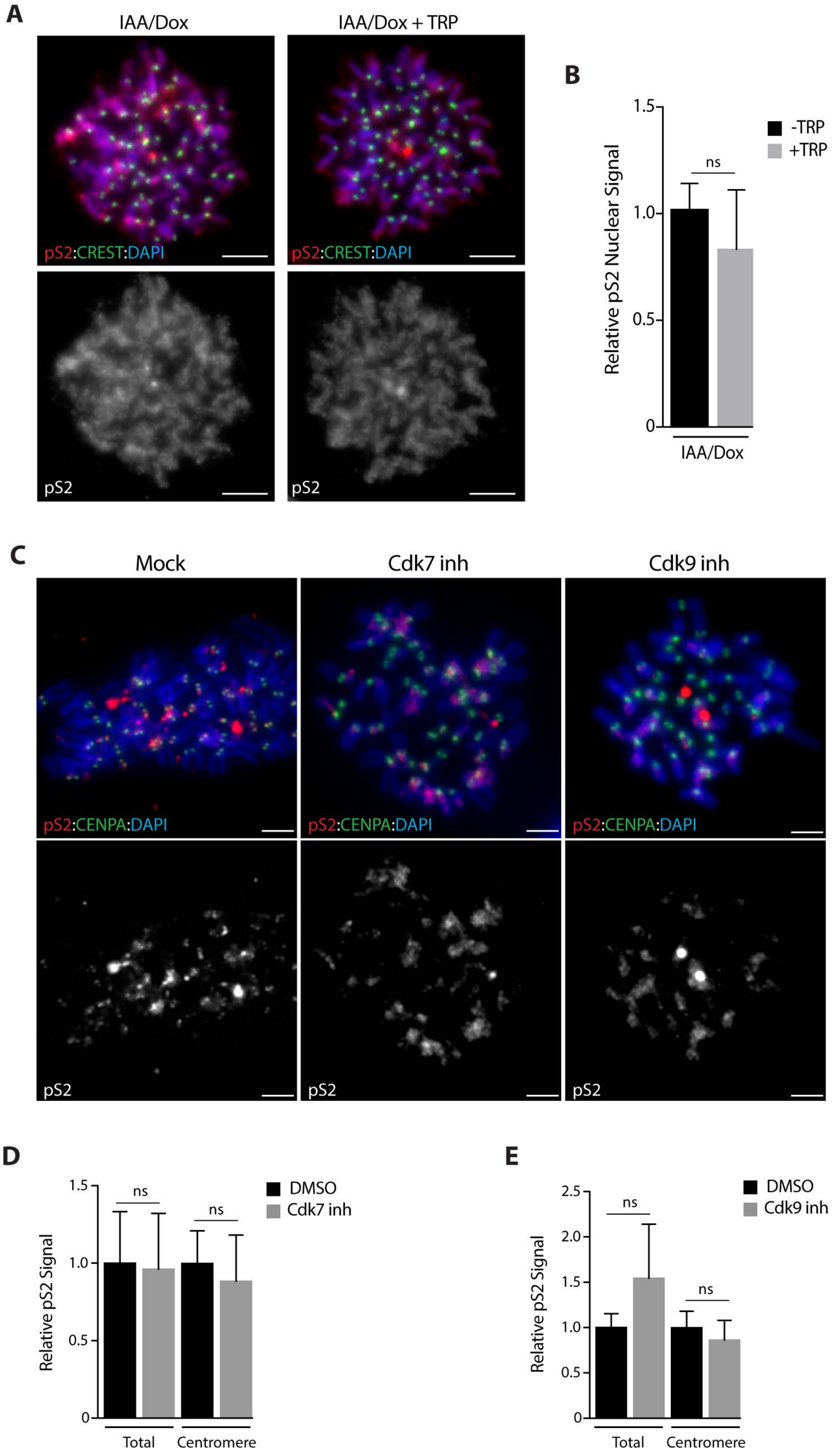


**Supplemental Figure 10**

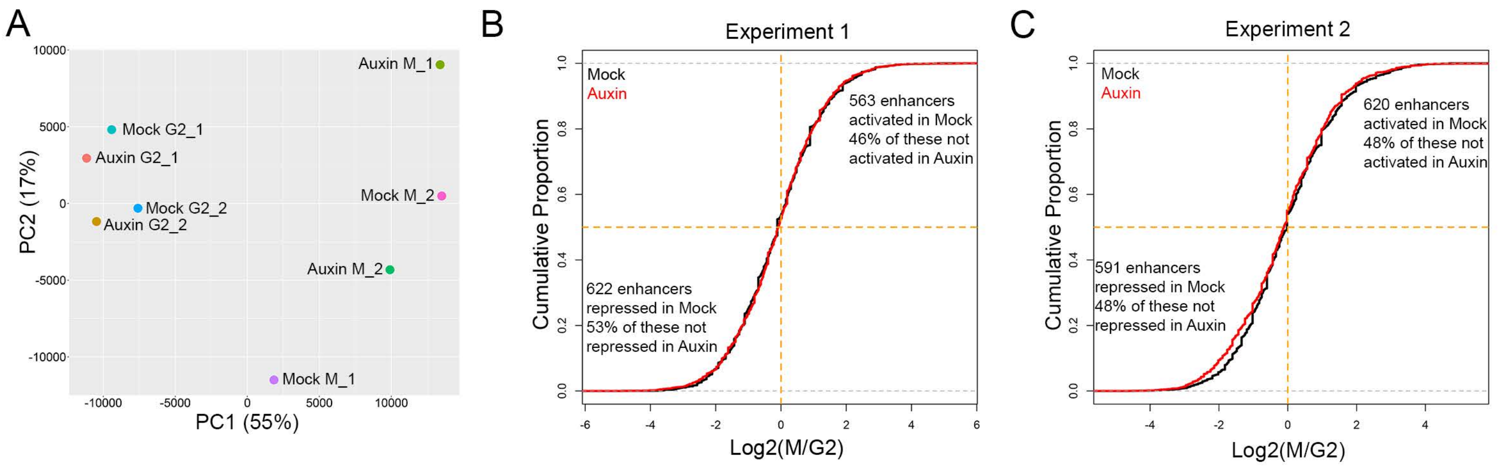


**Supplemental Figure 11.**

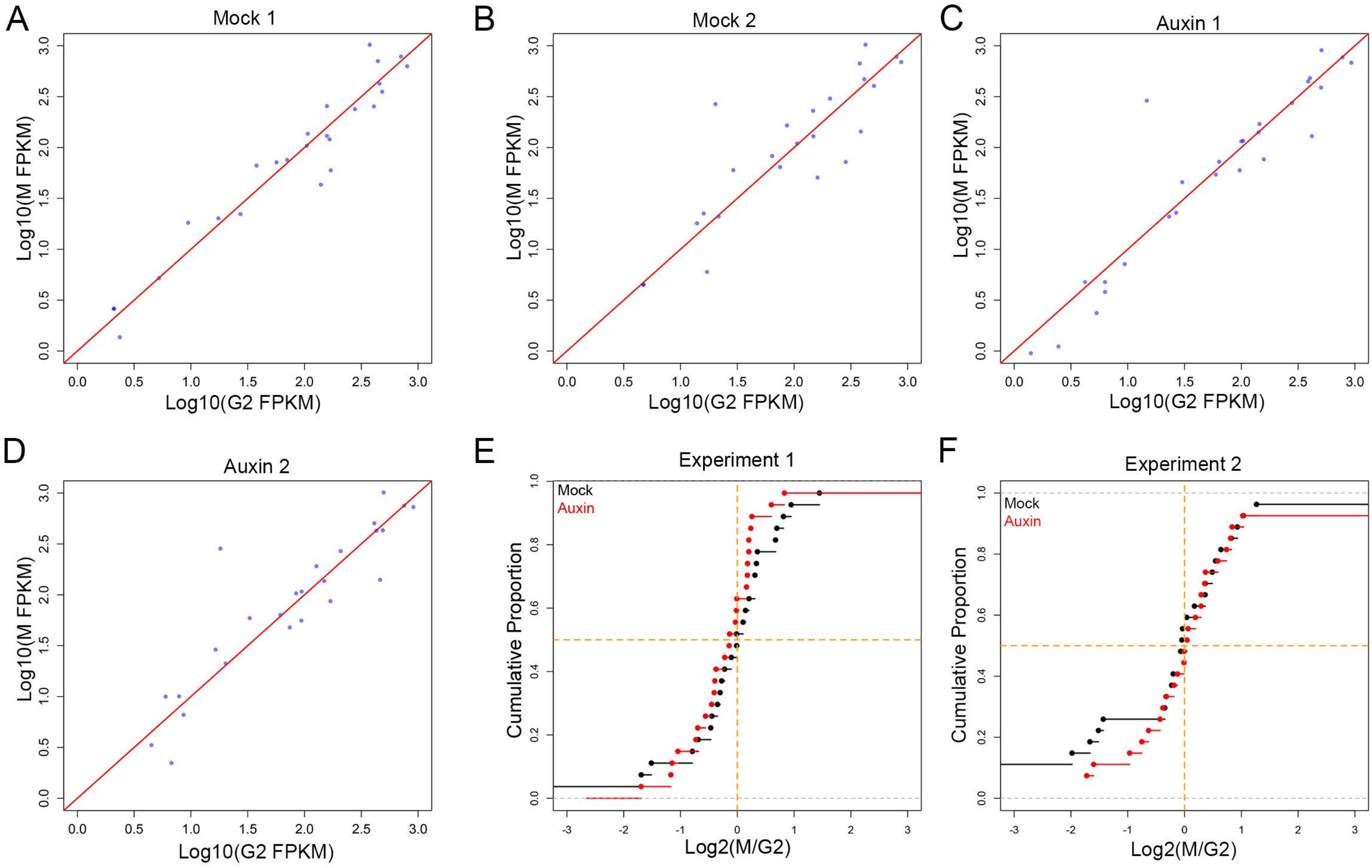


**Supplemental Figure 12**

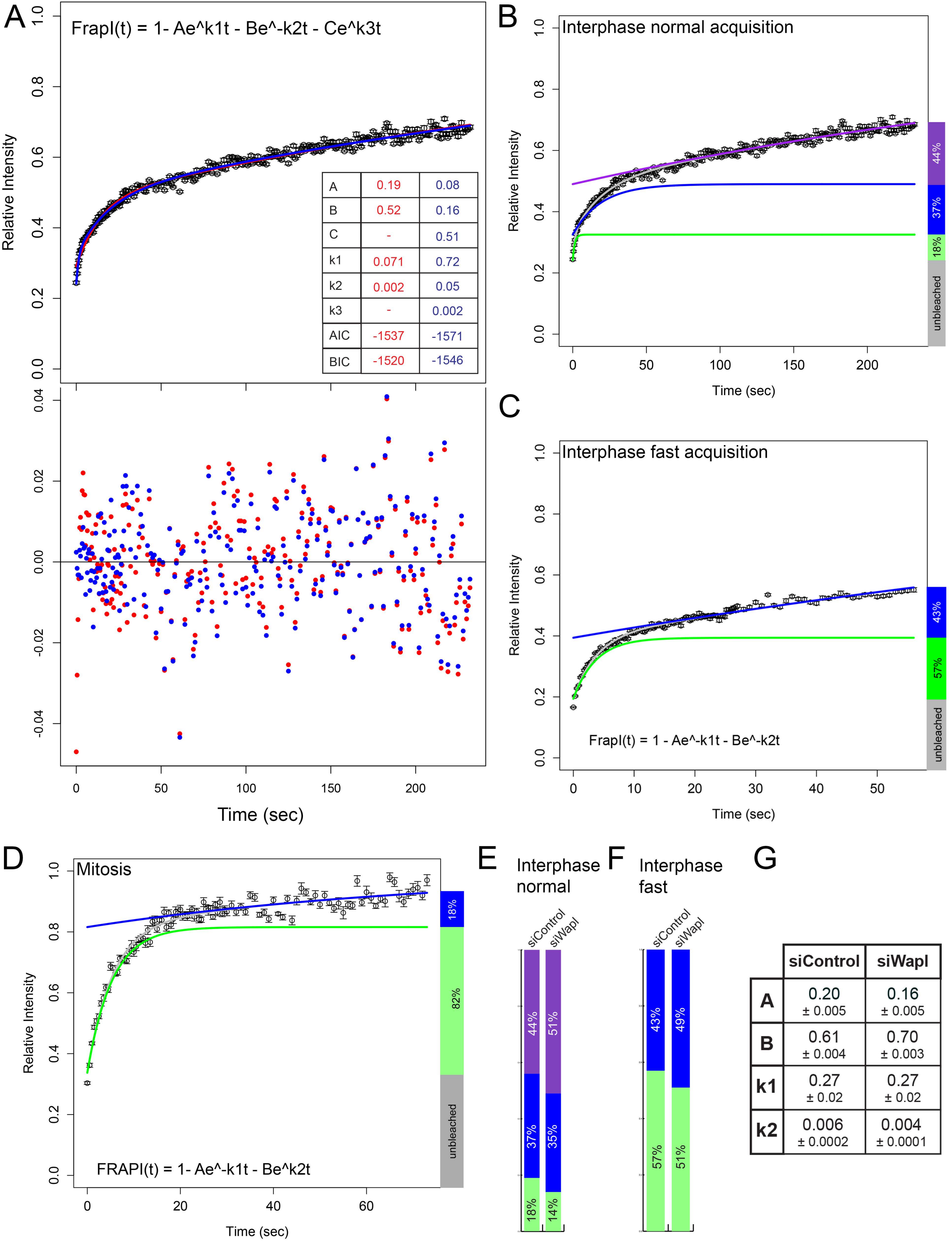


| | siControl | siWapl |
|---|---|---|
| A | 0.20 ± 0.005 | 0.16 ± 0.005 |
| B | 0.61 ± 0.004 | 0.70 ± 0.003 |
| k1 | 0.27 ± 0.02 | 0.27 ± 0.02 |
| k2 | 0.006 ± 0.0002 | 0.004 ± 0.0001 |

**Supplemental Figure 13**

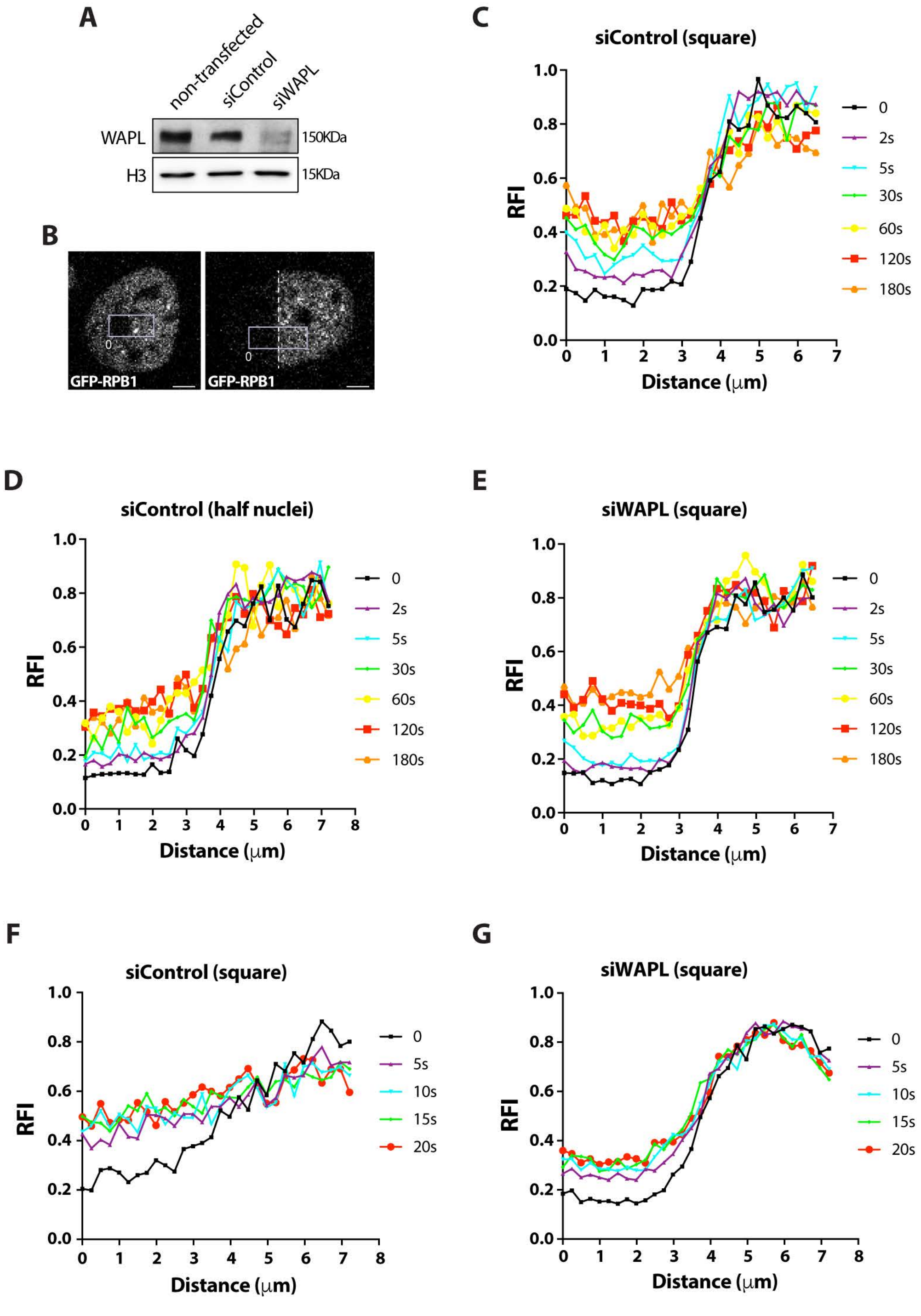


**Supplemental Figure 14**
